# Slow Gompertzian aging in long-lived *C. elegans* results from expansion of decrepitude, not decelerated aging

**DOI:** 10.1101/2025.04.13.648378

**Authors:** Bruce Zhang, David Gems

## Abstract

The Gompertz equation describes exponential age-increases in animal mortality rate arising from biological aging. Its parameters, *α* and *β*, are widely used to evaluate lifespan-extending interventions and human mortality patterns: it is assumed that reduction in *β* corresponds to deceleration of aging rate, and reduction in *α* to reduced aging-independent mortality. However, this view has never been empirically validated. We therefore investigated the biological basis of *α* and *β*, by simultaneous quantification of mortality and age-related health in long-lived populations of the nematode *Caenorhabditis elegans*. We show that reduction in *β* arises not from decelerated aging but expansion of decrepitude in longer-lived individuals, whereas reduction in *α* arises from decelerated aging. This empirical re-evaluation of Gompertzian aging inverts and challenges long-standing ideas in the biodemography of aging.

## Introduction

Aging (senescence) is the main cause of death and disease in the world today (Guo et al., 2022). Studies of the biology of aging often employ short-lived animal models such as the nematode *C. elegans*. Though biological senescence affects individuals, researchers investigating it frequently use demographic (population) senescence as a metric (Klass, 1977, Kenyon et al., 1993). However, demographic metrics (e.g. changes in mean lifespan) are difficult to link to determinative biological mechanisms. This is true of a salient and enigmatic feature of demographic aging: the exponential increase in mortality rate during adulthood, that is common (though not universal) among animal species (Jones et al., 2014, Finch et al., 1990). This was described in human populations two centuries ago by Benjamin Gompertz, and his eponymous equation *μ(x)=αe*^*βx*^ (Gompertz, 1825, Greenwood, 1928). Here, mortality rate *μ(x)* at age *x* is a function of a scale parameter *α* and rate parameter *β*, with the latter specifying the mortality rate acceleration with age. Accordingly, *α* and *β* together determine survival, mortality rate and death frequency functions (Fig. 1A).

**Fig. 1.**
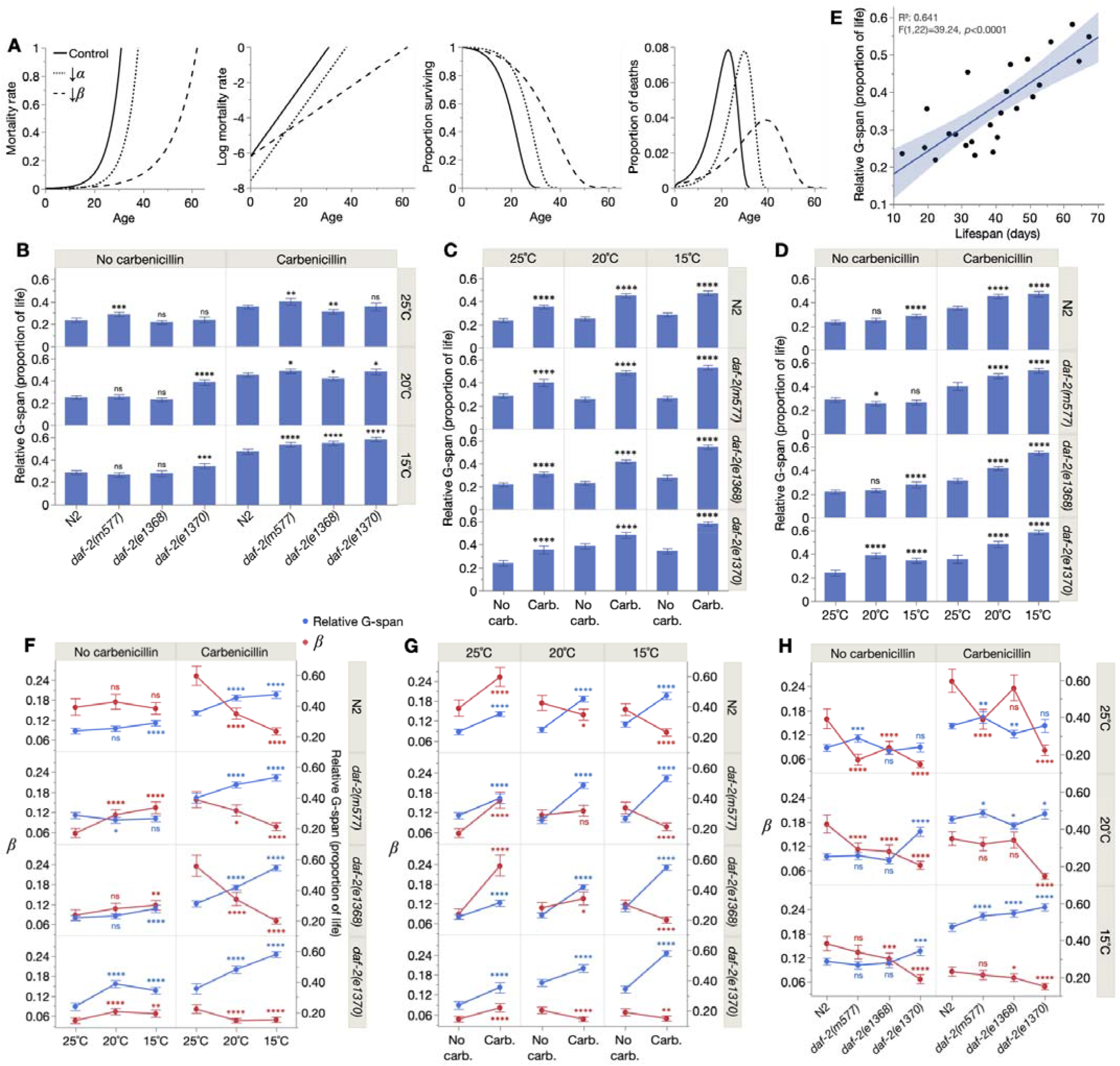
Effects of life-extending treatments on biological and demographic parameters. (**A**) Effects of decreasing *α* and *β* on demographic measures. Control: *α, β*: 0.002, 0.2; ↓*α*: *α, β*: 0.0005, 0.2; ↓*β*: *α, β*: 0.002, 0.1. First panel, effect on mortality rate. Here, *α* and *β* respectively control the scale and rate of exponential mortality rate increase. Second panel, effect on ln-transformed mortality rate. This transforms the exponential into a straight line, where *α* and *β* respectively control its intercept and gradient. Though a simple means to visualize *α* and *β* effects, this practice can be misleading and lead to improper parameter estimation (Eakin et al., 1995, Mueller et al., 1995, Shouman and Witten, 1995, Rozing and Westendorp, 2008). Third panel, effect on survival proportion. Here, *α* and *β* effectively shift and stretch, respectively, the survival curve along the x-axis. Fourth panel, effect on mortality frequency. Here, *α* and *β* effectively shift and stretch (respectively) the mortality frequency distribution along the x-axis. That is, reducing *β* (but not *α*) increases inter-individual lifespan variation. (B-E), Life-extending interventions disproportionately extend gerospan (G-span). Effects of (**B**) *daf-2(rf)*, (**C**) carbenicillin, and (**D**) reduced temperature on mean relative gerospan (G-span^rel^). *daf-2(e1368)* was previously observed to increase G-span^rel^ at 20°C (Podshivalova et al., 2017); that we did not see this could reflect differences in how G-span was measured. N2, wild-type. Statistical significance of G-span^rel^ differences were assessed by two-tailed Student’s t-tests, showing 95% confidence intervals; ns *p* > 0.05, * *p* ≤ 0.05, ** *p* ≤ 0.01, *** *p* ≤ 0.001, **** *p* ≤ 0.0001. (**E**) Mean G-span^rel^ is positively related to mean lifespan across the 24 cohorts, assessed by least-squares linear regression and F-test, showing the 95% confidence region. (F-H) Correspondence between reduced *β* and increased relative gerospan. Effects of (**F**) reduced temperature, (**G**) carbenicillin, and (**H**) *daf-2(rf)* on *β* and relative gerospan (G-span^rel^). N2, wild-type. G-span^rel^ values, 95% confidence intervals and statistical significance symbols are replotted from (B-D), and *β* calculated by maximum likelihood estimation, and statistical significance of *β* differences assessed by likelihood ratio tests, showing 95% confidence intervals.

But to what do these demographic parameters correspond in terms of biological aging, within individuals? Because *α* quantifies the initial mortality rate prior to aging (*x*=0, usually defined as in early adulthood), it is widely interpreted as relating to aging-independent mortality, arising e.g. from extrinsic hazards or intrinsic vulnerabilities (Sacher, 1977, Vaupel et al., 1979). By contrast, *β*, which corresponds to the exponential mortality rate increase (demographic senescence), is routinely taken to reflect biological aging rate. However, these interpretations remain conjectural and uncertain (Driver, 2001, Yashin et al., 2002, Driver, 2003, Masoro, 2006, Rozing and Westendorp, 2008, Koopman et al., 2011, Bansal et al., 2015, Hughes and Hekimi, 2016). A notable feature of the demography of aging is the high degree of inter-individual variability in lifespan. This is true of *C. elegans*, despite their being isogenic and maintained under identical conditions (Vaupel et al., 1998, Kirkwood et al., 2005). Inter-individual variation in age decline and lifespan in *C. elegans* has been investigated in several previous studies (Huang et al., 2004, Hahm et al., 2015, Stroustrup et al., 2016, Zhang et al., 2016, Churgin et al., 2017, Zhao et al., 2017, Newell Stamper et al., 2018, Statzer et al., 2022). Given that *β* is a function of inter-individual variation in lifespan (Fig. 1A, last panel), characterizing the latter is key to understanding this parameter.

Here we explore the biological meaning of *α* and *β* in terms of age changes at the individual level. To this end, we simultaneously quantified biological and demographic aging by means of longitudinal analysis of numerous individual *C. elegans* subjected to different life-extending interventions. In this analysis, *β* proved to be a measure of variability in gerospan (late-life decrepitude) rather than biological aging rate, and *α* more of biological aging rate than of aging-independent processes.

## Results

### Three life-extending interventions disproportionately extend gerospan

We examined the effects on biological and demographic aging of three types of intervention that increase *C. elegans* lifespan: (i) reduction of ambient temperature, from 25°C to 20°C or 15°C (Klass, 1977), (ii) prevention of late-life infection (by dietary *E. coli*) using antibiotics (Garigan et al., 2002), and (iii) reduction-of-function (*rf*) mutation of the *daf-2* insulin/IGF-1 receptor (Kenyon et al., 1993), specifically the class 1 alleles *daf-2(m577)* and *daf-2(e1368)*, and class 2 (more pleiotropic) allele *daf-2(e1370)* (Gems et al., 1998).

Lifespan was measured for all 24 combinations of conditions (3 temperatures, ± carbenicillin, 4 genotypes) in 6 successive trials. To relate the Gompertz parameters to individual-level biological aging features, in 4/6 trials, Gompertz parameters were estimated and animals tracked individually and locomotory health assessed throughout life (Hosono et al., 1980, Herndon et al., 2002). Animals were scored as either youthful (healthy, sinusoidal locomotion) or decrepit (non-sinusoidal/uncoordinated locomotion, or immotility). We will refer to absolute and relative healthspan (H-span^abs^, H-span^rel^) and gerospan (G-span^abs^, G-span^rel^) to describe, respectively, the number of days or proportion of life spent in youthfulness or decrepitude (morbidity). Necropsies were also performed to determine the mode of death of all animals (Zhao et al., 2017).

All lower temperature, antibiotic and *daf-2(rf)* treatments increased mean lifespan (average effects: +42.0%, +58.9%, +77.1%, respectively; fig. S1, table S1), as expected (Klass, 1977, Kenyon et al., 1993, Garigan et al., 2002). We first examined effects of life-extending treatments on G-span^rel^. It was previously observed that on infective *E. coli*, G-span^rel^ is increased by *daf-2(rf)* (Huang et al., 2004, Bansal et al., 2015, Podshivalova et al., 2017) and, notably, that this increase is diminished if infection is prevented, due to disproportionately increased survival of decrepit wild-type individuals (Podshivalova et al., 2017, Statzer et al., 2022). Thus, greater G-span^rel^ in *daf-2(rf)* mutants appears to reflect their resistance to infection.

At 20°C, *daf-2(e1370)* increased G-span^rel^ (relative to N2), and this increase was largely suppressed by carbenicillin (Fig. 1B), consistent with infection resistance unmasking decrepitude (Podshivalova et al., 2017). However, among other *daf-2(rf)* allele/temperature combinations, G-span^rel^ patterns varied markedly (Fig. 1B); thus, *daf-2(rf)* has allele- and condition-specific effects on G-span^rel^. However, only 2/18 *daf-2(rf)* treatments decreased G-span^rel^ (relative to N2) compared to 9/18 that increased it, i.e. longevity usually entailed expanded decrepitude. In addition, 12/12 carbenicillin and 12/16 lower temperature treatments also increased G-span^rel^ (Fig. 1C-D). Direct regression of G-span^rel^ against lifespan for the 24 cohorts revealed a strong positive relationship (Fig. 1E). Thus, the longevity interventions tested generally increase the proportion of life in poor health, and do not cause a simple deceleration of aging (which should stretch H-span^abs^ and G-span^abs^ proportionally, leaving G-span^rel^ unchanged).

### Correspondence between reduced *β* and increased relative gerospan

Next, we considered the central question of this study: the relationship between biological and demographic aging. To this end, the Gompertz parameters *α* and *β* were obtained for the 24 cohorts by maximum likelihood estimation (Pletcher, 1999), and relationships between life-extending treatment effects on the parameters and G-span^rel^ examined; we start with *β*, reduction of which has often been equated with decelerated biological aging.

First, effects of reducing temperature. On carbenicillin, this decreased *β* in all genotypes (Fig. 1F), which would be consistent with a general slowing of the rate of living. However, without carbenicillin, *β* was unchanged in wild-type and even increased in *daf-2* mutants, suggesting complex effects of genotype and temperature on host-*E. coli* interactions. Surprisingly, *β* reduction by low temperature (on carbenicillin) consistently co-occurred with G-span^rel^ increase (Fig. 1F, right), suggesting that *β* reduction here reflects neither simply slowed nor healthier biological aging.

Next, antibiotic treatment. Against the expectation that preventing infection (a largely extrinsic insult) would reduce *α* alone, carbenicillin decreased *β* in all genotypes at 15°C, and in wild-type and *daf-2(e1370)* at 20°C (Fig. 1G). In each case, G-span^rel^ was again increased. In the remaining carbenicillin treatments, both *β* and G-span^rel^ increased, suggesting that *β* reduction may require G-span^rel^ increase, but not vice versa.

Third, *daf-2(rf)* mutation. *daf-2(e1370)*, the severest (and longest-lived) allele, also lowered *β* while increasing G-span^rel^, both off and on carbenicillin, at 15°C and 20°C (Fig. 1H). At 25°C, *β* was decreased without change in G-span^rel^, an exception to the pattern; similarly, only 3/7 class 1 *daf-2(rf)* treatments reducing *β* increased G-span^rel^. Despite these exceptions, 21/27 (78%) of all treatments that significantly reduced *β* increased G-span^rel^. Thus, amongst these life-extending interventions, reduction of *β* reflects expanded decrepitude rather than a simple deceleration of biological aging. Of course, this does not preclude more complex forms of aging deceleration, as demonstrated further below.

### Reduced *β* reflects inter-individually variable gerospan expansion

Given that *β* is a function of lifespan variation (Fig. 1A, fourth panel), we wondered about the relation of *β* to inter-individual differences in aging. As expected, amongst our 24 treatments, *β* showed strong inverse correlation with lifespan standard deviation (Fig. 2A). According to a previous estimate, G-span differences account for most (∼67%) inter-individual lifespan variation in normal-lived *C. elegans* (Zhang et al., 2016). Thus, the correspondence between reduced *β* and increased G-span^rel^ in our cohorts might reflect inter-individually variable G-span expansion. To investigate this, we assessed H-span^abs^ and G-span^abs^ for all individuals in the 21 treatments displaying an inverse *β*-G-span^rel^ relationship (Fig. 1F-H).

**Fig. 2.**
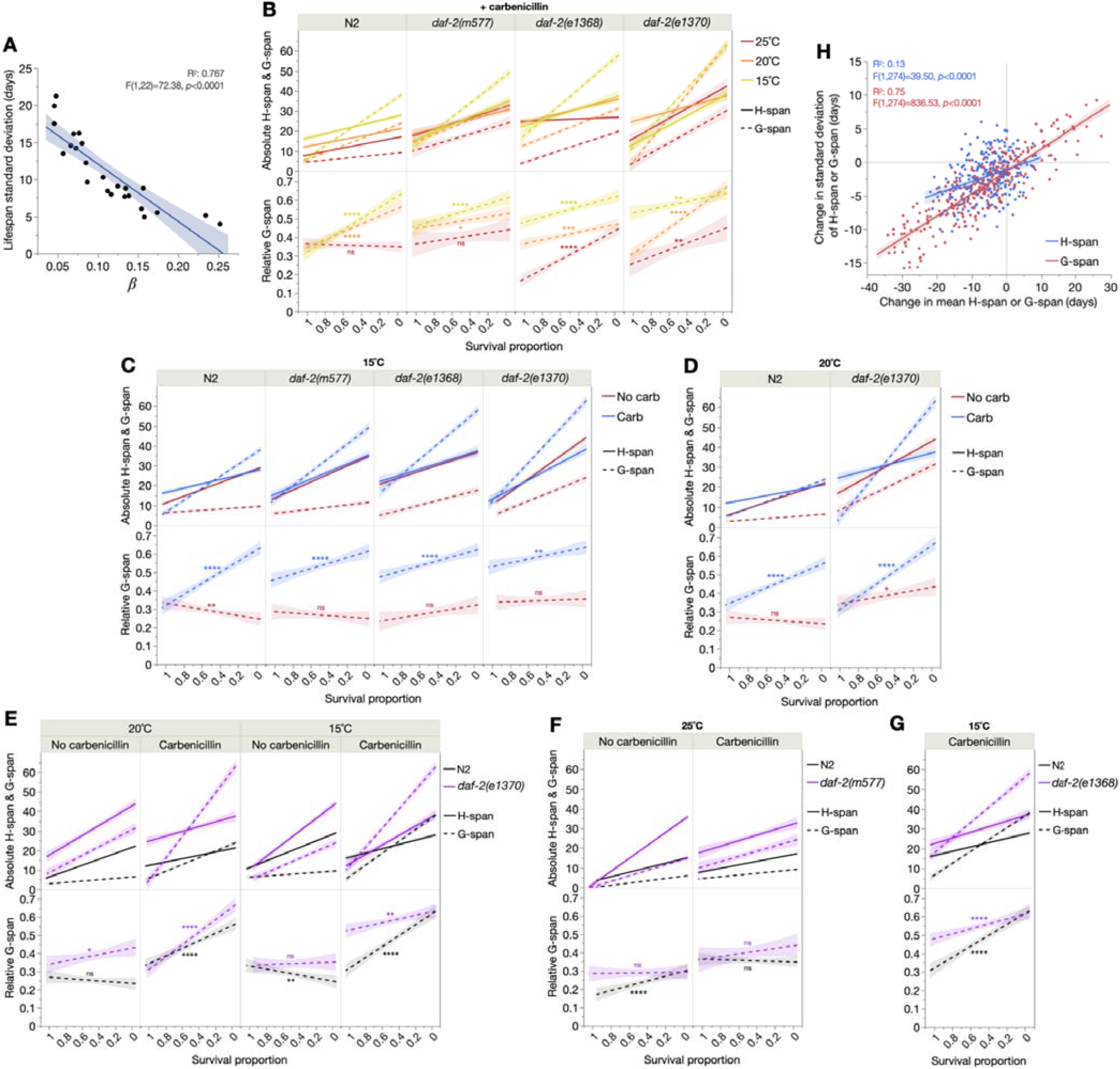
Reduced *β* reflects inter-individually variable gerospan expansion. (**A**) *β* is an inverse measure of lifespan standard deviation, across the 24 cohorts, assessed by least-squares linear regression and F-test, showing the 95% confidence region. Effects of (**B**) low temperature, (**C**-**D**) carbenicillin, and (**E**-**G**) *daf-2(rf)* on absolute healthspan and gerospan (upper panels) and relative gerospan (lower panels), plotted over survival proportion (i.e. x-axis left: shorter-lived individuals, x-axis right: longer-lived individuals). All panels show least-squares linear regressions with 95% confidence regions, assessed by F-tests. (**H**) Expansion of G-span is a more inter-individually variable process than expansion of H-span, across all possible pairs of the 24 cohorts. In addition to a higher R^2^, the steeper G-span linear fit indicates that G-span variation increases faster than H-span variation under life-extending conditions. Consistent with this, the range of values for mean H-span and H-span standard deviation is less than that for G-span. These least-squares linear regressions were assessed by F-tests, showing the 95% confidence region.

On carbenicillin, reducing temperature to 20°C in wild-type, *daf-2(m577)* and *daf-2(e1370)* increased G-span^abs^ disproportionately more than H-span^abs^ in longer-lived individuals (Fig. 2B top, fig. S2A top and middle). As a result, G-span^rel^ was increased in longer-lived individuals (Fig. 2B bottom, fig. S2A bottom). This illustrates how life-extending interventions can act by amplifying the “extended twilight” (Zhang et al., 2016) (greater G-span^rel^) of longer-lived individuals within a population, which we will refer to as *extended twilight longevity* (ETL). Notably, in ETL, *β* reduction and the associated extension of the survival curve tail reflects the disproportionate expansion of decrepitude in longer-lived individuals. Meanwhile in *daf-2(e1368)* and 15°C in *daf-2(e1370)*, G-span^rel^ increased more in *shorter*-lived individuals, resulting in an inverted ETL (Fig. 2B).

In all carbenicillin treatments yielding the inverse *β*-G-span^rel^ relationship, G-span^abs^ again increased disproportionately more than H-span^abs^ in longer-lived individuals, thus increasing G-span^rel^ in these individuals (Fig. 2C-D, fig. S2B). Thus, carbenicillin too is an ETL treatment that lowers *β* via variable G-span expansion; this is particularly striking, since preventing infection is not generally expected to affect biological aging rate, unlike temperature.

Finally, we asked if *β* reduction in *daf-2(rf)* reflects ETL. At 20°C, and 15°C without carbenicillin, *daf-2(e1370)* indeed increased G-span^abs^ more than H-span^abs^, and therefore G-span^rel^, in longer-lived individuals (Fig. 2E, fig. S2C). However, on carbenicillin at 15°C, *daf-2(e1370)* showed inverted ETL. Of the three class 1 *daf-2(rf)* treatments with an inverse *β*-G-span^rel^ relationship, one exhibited modest ETL, and two inverted ETL (Fig. 2F-G, fig. S2C).

In summary, *β* reduction reflected ETL in 5/8 reduced temperature, 6/6 antibiotic, and 4/7 *daf-2(rf)* treatments (15/21 or 71%, in total) where *β* reduction corresponded to G-span^rel^ increase. Notably, in all 6 exceptions (showing inverted ETL), control cohorts already exhibited extended twilight in long-lived individuals (Fig. 2B, E-G), perhaps limiting additional ETL.

As a further probe of whether *β* reduction arises from ETL in these 15 treatments, we compared contributions of H-span^abs^ and G-span^abs^ changes to *β* reduction, and to lifespan variation increase (i.e. *β* reduction). Indeed, G-span^abs^ changes decreased *β* more than H-span^abs^ changes in 13/15 treatments (compared to 0/6 exceptions with inverted ETL) (table S2), and increased lifespan variation more in 14/15 interventions (compared to 3/6 exceptions) (table S3).

Consistent with these findings, across the 24 cohorts G-span variation increased more rapidly with mean G-span than H-span variation with mean H-span, while mean length and variability of G-span had wider ranges than that of H-span (Fig. 2H). This shows that across these cohorts, inter-individually variable G-span expansion (i.e. ETL), rather than H-span expansion, is the predominant mode of life extension.

These findings argue that expanded decrepitude not only correlates with a lower *β*, but causes it. That is, gerospan increase in longer-lived individuals causes the extended survival curve tail that is characteristic of reduced *β* (Fig. 1A, third panel). Therefore, *β* here is not a measure of biological aging rate but rather of inter-individual heterogeneity in late-life decrepitude.

### Reduced *α* reflects slowed biological aging rate

Because healthspan expansion is generally considered a reliable indicator of slowed biological aging (given more time to G-span onset), we wondered if *β* reduction could at least indirectly reflect decelerated aging, should G-span^abs^ and H-span^abs^ expansions occur together. Indeed, H-span^abs^ was increased in 9/15 ETL treatments (Fig. 3A-C). We therefore examined the effect of H-span^abs^ expansion on the Gompertz parameters in these 9 treatments. H-span^abs^ expansion significantly decreased *β* in 4/9 treatments but, strikingly, significantly decreased *α* in 8/9 (Fig. 3D, table S4A-C). This suggests that *α*, though not traditionally associated with biological aging, may in fact be a better measure of it than *β*.

**Fig. 3.**
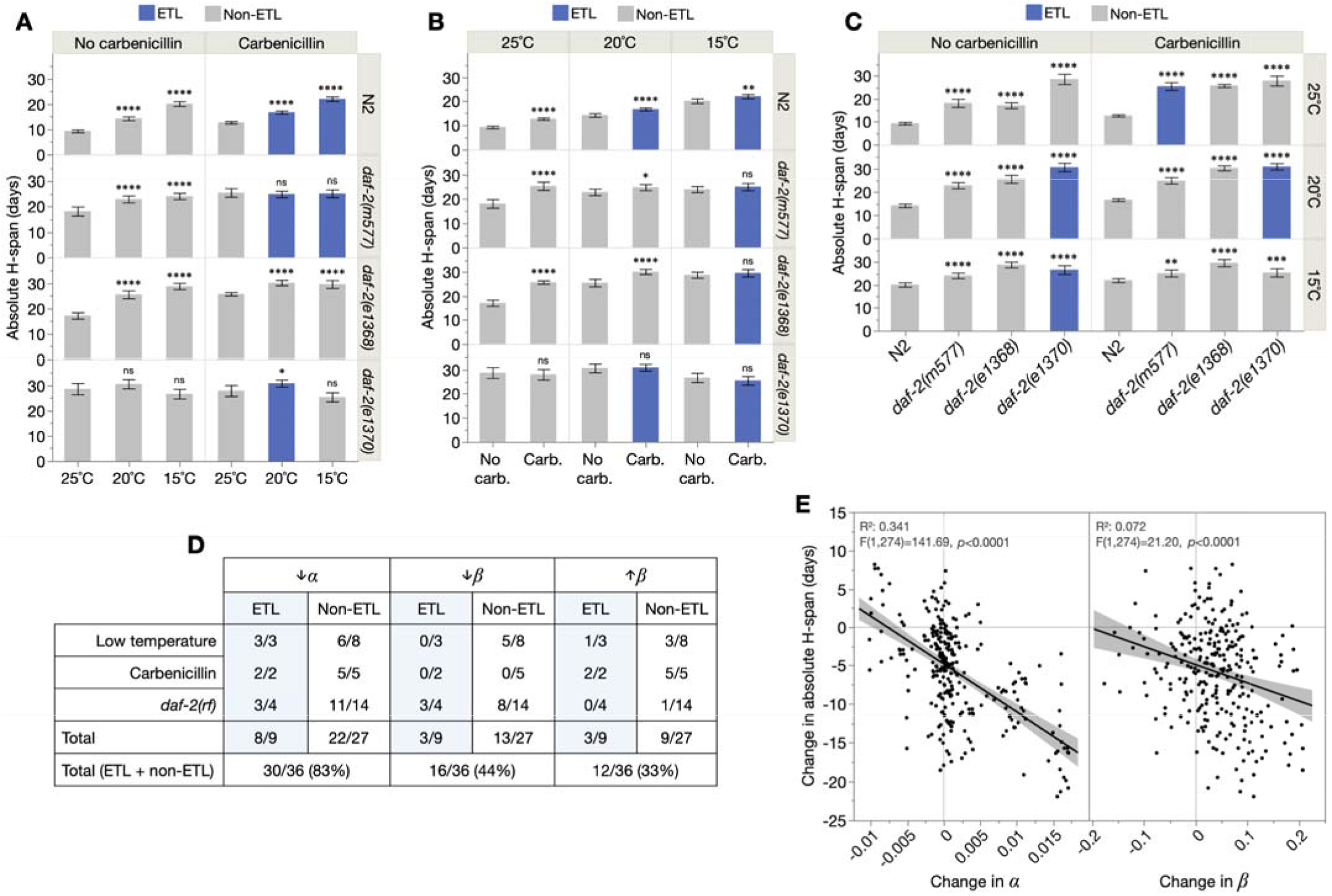
*α* outperforms *β* as a measure of absolute healthspan. Effects of (**A**) low temperature, (**B**) carbenicillin and (**C**) *daf-2(rf)* on mean absolute healthspan (H-span^abs^). N2, wild-type. ETL, extended twilight longevity. H-span^abs^ differences were assessed by two-tailed Student’s t-tests, showing 95% confidence intervals. (**D**) Table summarizing fractions of ETL and non-ETL treatments in which H-span^abs^ increase causes the specified Gompertz parameter change, for each life-extending treatment class. (**E**) Least-squares linear regressions of the changes in H-span^abs^ between all possible pairs of the 24 cohorts, over the corresponding change in *α* or *β* for those pairs. This shows that *α* is the better predictor of H-span^abs^. The relationships were assessed by F-tests, showing the 95% confidence region.

To assess if this unexpected finding is idiosyncratic to ETL, we also examined the non-ETL treatments, of which 27/31 significantly extended H-span^abs^ (Fig. 3A-C). However, again, H-span^abs^ expansion decreased *α* in 22/27 (81%) of these non-ETL treatments, but decreased *β* in only 13/27 (48%) (Fig. 3D, table S4A-C). Additionally, H-span^abs^ expansion *increased β* in 9/27 treatments, including in all carbenicillin treatments; thus, slowing biological aging can even increase *β*. Considering all treatments (ETL plus non-ETL), H-span^abs^ expansion decreased *α* in 30/36 (83%) cases, decreased *β* in 16/36 (44%), and increased *β* in 12/36 (33%). This suggests that *α* should better predict H-span^abs^ than *β* amongst the 24 cohorts, and this indeed proved to be the case (Fig. 3E). In summary, healthspan expansion, a plausible metric of slowed biological aging, more consistently reduces *α* than *β*, and even increases *β*. Thus, overall, our empirically-based findings invert traditional interpretations of the biological meaning of the two Gompertz parameters.

### *α* and *β* describe inter-individual heterogeneity in age-related bacterial pathology

To further understand the biological mechanisms underpinning the Gompertz parameters, we performed necropsies on all corpses, building upon established mortality deconvolution methodology (Zhao et al., 2017). We scored for aging-related bacterial colonization of the pharynx and intestine (Fig. 4A, fig. S3A-B; no carbenicillin). We noted that almost all corpses with a swollen, infected pharynx (P or “big P”, as opposed to p or “small p” with an atrophied, uninfected pharynx) (Zhao et al., 2017) also had intestinal colonization (IC) by *E. coli*, but not vice versa. Thus, P animals are largely a subset of those with IC (fig. S3). Death type was accordingly used to define three biologically-distinct subpopulations: P (i.e. PIC), pIC (<u>p</u> with <u>IC</u>), and pnIC (<u>p</u> with <u>n</u>o <u>IC</u>). Consistent with prior findings (Zhao et al., 2017), lifespan was shorter for P than p (pIC and pnIC) subpopulations for most treatments (Fig. 4B, table S5). Lifespan was also shorter for pIC than pnIC populations (at 15°C and 25°C, but not 20°C). Thus, the three subpopulations exhibit distinct aging trajectories.

**Fig. 4.**
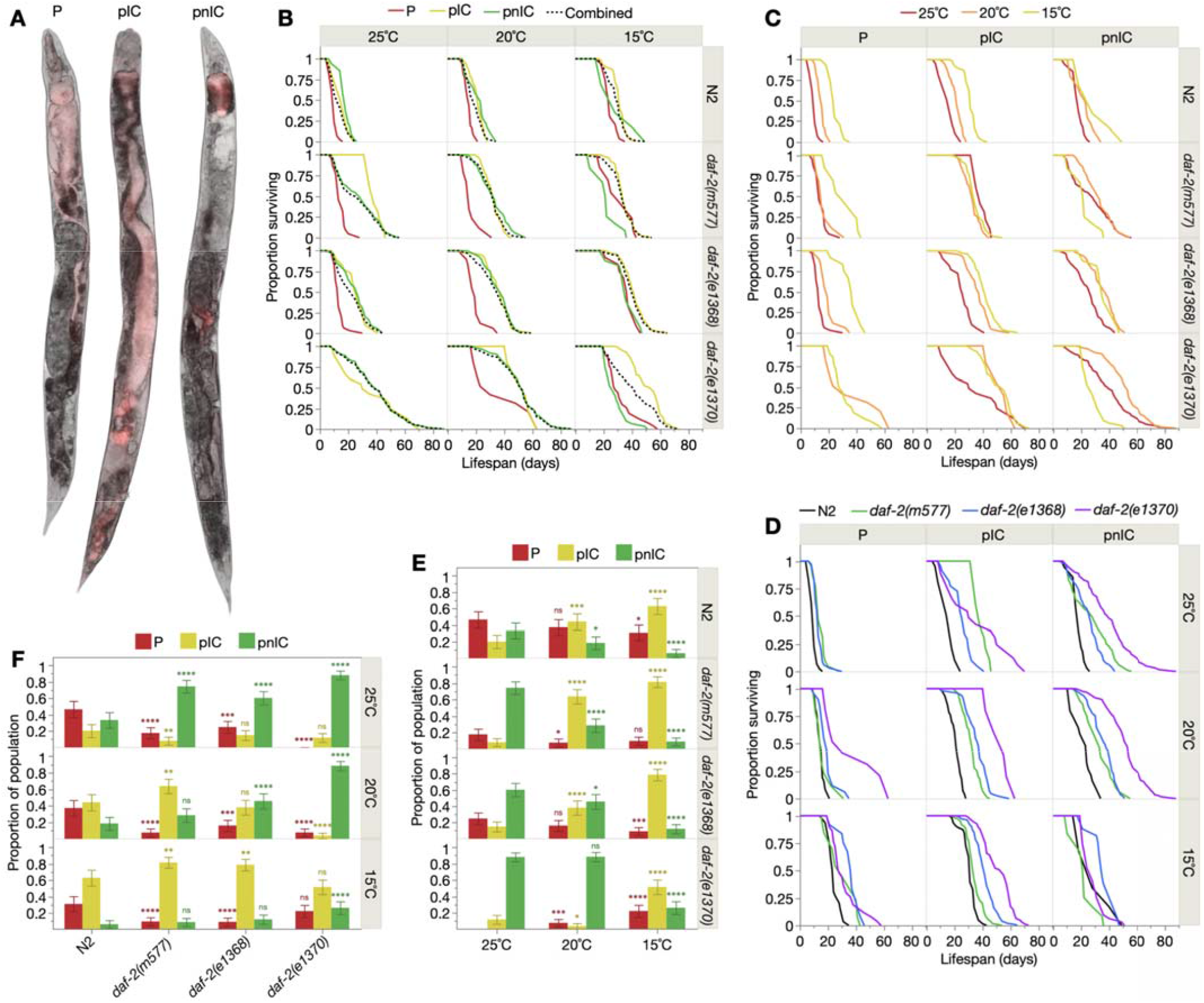
Life-extending interventions change lifespan and prevalence of pathological subpopulations. (**A**) Images of representative P, pII, and pnII corpses, fed throughout life with RFP-expressing *E. coli*; overlay of brightfield and epifluorescence images. (**B**) Survival curves of P, pIC and pnIC subpopulations and whole population (“Combined”). (C-D) Effect of (**C**) low temperature and (**D**) *daf-2(rf)* treatments on subpopulation lifespan. Log-Rank *p* values of depicted comparisons for B-D are presented in table S5. Survival proportions were obtained from Kaplan-Meier lifespan analysis using pseudofrequencies (in place of frequency) to account for censors (see Methods for details). (E-F) Effects of (**E**) low temperature and (**F**) *daf-2(rf)* on subpopulation prevalence. Differences in prevalence was assessed by Pearson’s chi-squared test.

We then investigated how inter-subpopulation heterogeneity affects *β*. Notably, *β* was consistently smaller in whole populations than in subpopulations (table S6). This illustrates how *β* can underestimate demographic aging rate in the presence of hidden subpopulation heterogeneity (Yashin et al., 2002), and according to the traditional interpretation of *β*, paradoxically predicts that subpopulations age biologically faster than their combination (whole population). This underscores how *β* can be a measure of inter-individual heterogeneity in aging, rather than of biological aging rate itself.

We next examined how life-extending interventions (here low temperature and *daf-2(rf)*, without carbenicillin) affect subpopulation prevalence and lifespan. While both treatments increased lifespan in most subpopulations, increases were generally greater for P by low temperature, and pIC by *daf-2(rf)* (Fig. 4C-D; table S5). Lower temperature modestly decreased P prevalence in all genotypes except *daf-2(e1370)*, suggesting reduced bacterial pathogenicity and/or enhanced host immunity, and allele-specific temperature sensitivity in *daf-2(e1370)* (Zhao et al., 2021) (Fig. 4E). Interestingly, however, low temperature strongly increased pIC prevalence (concomitantly decreasing pnIC prevalence). Thus, reducing temperature may act antagonistically on terminal infection, respectively decreasing and increasing pharyngeal and intestinal colonization. Meanwhile, *daf-2(rf)* treatment also decreased P prevalence, as previously seen (Zhao et al., 2021) and affected pIC prevalence in an allele- and temperature-specific manner (Fig. 4F), largely consistent with known *daf-2(rf)* infection resistance (Podshivalova et al., 2017).

We next determined the relative contribution of these changes in subpopulation lifespan and prevalence to the Gompertz parameters. Effects of low temperature on *β* proved to be determined mainly by changes in pnIC lifespan, followed closely by the increase in pIC to pnIC ratio in *daf-2(m577)* (Fig. 5A, fig. S4A). In *daf-2(rf)* treatment, changes in *β* were again primarily determined by changes in pnIC and/or pIC lifespan, rather than in P lifespan or subpopulation prevalence (Fig. 5B, fig. S4B). Thus, *β* is largely a function of p lifespan; specifically, increases and decreases in *β* typically result, respectively, from decreased and increased p (pIC and/or pnIC) longevity. This further demonstrates that *β* is not a measure of biological aging rate but inter-individual heterogeneity, here manifested as subpopulation-specific responses to longevity interventions.

**Fig. 5.**
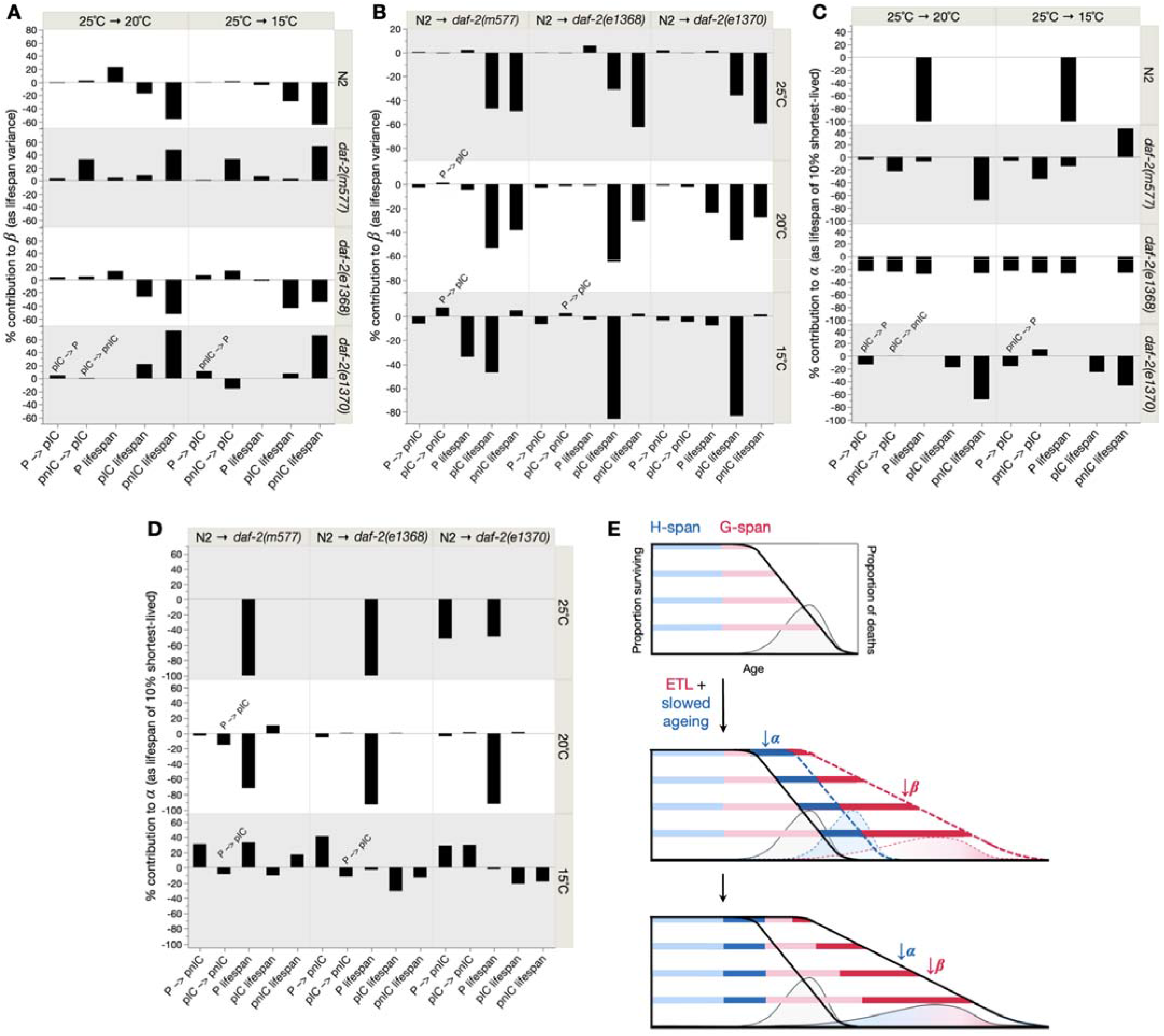
*α* and *β* reflect subpopulation-specific changes in lifespan and prevalence. Relative contributions (sum to 100%) of changes in subpopulation prevalence and lifespan to overall change in population (**A**-**B**) *α* and (**C**-**D**) *β* in (A,C) low temperature and (B,D) *daf-2(rf)* treatments. Bar height indicates contribution magnitude (%) and bar direction (+ or –) indicates contribution direction (increase or decrease Gompertz parameter). The first two x-axis items are component contributions of changes in subpopulation prevalence (based on principles of parsimony, Fig. 4E-F), while the remaining x-axis items are contributions of changes in subpopulation lifespan. Overlaid angled labels reflect different subpopulation prevalence changes for those specific treatments. See Methods for full analysis details. (**E**) Summary schematic of the biological basis of the Gompertz parameters across our experimental cohorts, depicting a hypothetical intervention that reduces both *α* and *β*. Each panel depicts healthspan (H-span; blue bar segment) and gerospan (G-span, red bar segment) for four individuals representative of their depicted lifespan within the population (age at end of bar), as bordered by the survival curves (left y-axis, bold data lines). Here, as demonstrated in *C. elegans*, reduction in *α* arises from H-span expansion (equally across individuals in this simplified depiction), leading to the approximately parallel right-shift of the survival curve (middle panel, dashed blue) that occurs in reduction of *α*. In contrast, reduction in *β* arises from inter-individually variable G-span expansion that is greater in longer-lived individuals, leading to the approximate horizontal stretch of the survival curve (middle panel, dashed red) that occurs in reduction of *β*. The bottom panel shows the overall effect of this hypothetical intervention (reduces both *α* and *β*), with H-span and G-span arranged in order. In all panels, probability distributions of death times corresponding to the survival curves, are overlaid (right y-axis, thin data lines), showing that reduction in *α* causes an approximate right-shift of death ages, whereas reduction in *β* causes an additional stretch that increases lifespan variation. The area under these probability distributions is shaded to reflect the level of variation in H-span/G-span between individuals of different lifespans. This shows that reduction in *α* causes an approximate right-shift of death times, due to similar expansion of H-span in all individuals (solid blue shading), whereas reduction in *β* stretches the distribution of death times, due to greater G-span expansion in longer-lived individuals (red gradient shading). Therefore, *β* describes the degree of inter-individual variation in health and lifespan, and not biological aging rate, which is better captured by *α* as H-span length.

We similarly deconvolved determinants of *α*. In low temperature treatment, determinants of *α* were genotype-specific: *α* was wholly determined by P lifespan in wild-type, mainly by pnIC lifespan in *daf-2(m577)* and *daf-2(e1370)*, and by all determinants except pIC lifespan in *daf-2(e1368)* (Fig. 5C, fig. S4A). In *daf-2(rf)* treatment, *α* was primarily determined by P lifespan at 25°C and 20°C, while at 15°C, several determinants were important; in common, the increase in pnIC to P ratio (Fig. 5D, fig. S4B). Therefore, in contrast to *β, α* is largely a function of P lifespan and subpopulation prevalence, but like *β*, reflects subpopulation-specific responses to life-extending interventions. These findings illustrate how the Gompertz parameters can reflect biological changes affecting some but not all individuals, and are therefore unsuitable measures in heterogeneous (i.e. most, and particularly human) populations.

## Discussion

This study defines the biological underpinnings of Gompertzian demographic aging in *C. elegans* under 24 conditions. It demonstrates how the Gompertz rate parameter *β* corresponds to inter-individual gerospan variation rather than biological aging rate, while the Gompertz scale parameter *α* better reflects biological aging rate, rather than aging-independent determinants of mortality (Fig. 5E). Additionally, both parameters are the product of subpopulation-specificity in patterns of late-life disease. These findings underscore the great tenuousness of the correspondence between parameters describing population aging on the one hand, and the biological aging rate of component individuals on the other (Yashin et al., 2002).

We have shown how reduction of *β*, usually interpreted as slowed aging, can reflect gerospan expansion in longer-lived individuals, which stretches the survival curve tail and increases lifespan variation. Consequently, longer-lived individuals experience extended twilights (i.e. disproportionately-extended gerospans), as previously described (Zhang et al., 2016, Churgin et al., 2017, Statzer et al., 2022). Here we describe how this occurs *between* populations, so-called *extended twilight longevity* (ETL). Thus, here *β* reflects not biological aging rate, but gerospan variability in ETL (Fig. 5E). Such ETL could contribute to the temporal scaling of survival curves previously described in *C. elegans* (Stroustrup et al., 2016).

Given the gradual nature of aging, any delineation of healthspan and gerospan is inevitably somewhat arbitrary (Zhang et al., 2016, Newell Stamper et al., 2018). It is after the highly coordinated stages of development and reproduction that aging begins and consequent inter-individual variation in age decline emerges, in midlife by one estimate (Uno et al., 2025). Thus, it is likely that the contribution of gerospan variation to *β* has been underestimated, given that gerospan as we defined it excluded earlier ages with subtler senescent changes.

Regarding the Gompertz scale parameter *α*, past conjectures as to its biological meaning include intrinsic, aging-independent “vulnerability” or “frailty”, and extrinsic, age-related hazards, but not biological aging rate itself (Sacher, 1977, Finch et al., 1990). We have shown that *α* outperforms *β* as a predictor of healthspan length, arguing against the “not aging” interpretation, and providing experimental support for earlier, theoretical doubts in this regard (Masoro, 2006, Rozing and Westendorp, 2008, Blagosklonny, 2010) (Fig. 5E).

Are such nematode-derived reinterpretations of *α* and *β* likely to be applicable to higher animals? The frequent occurrence of Gompertzian aging throughout the animal kingdom (Finch et al., 1990, Jones et al., 2014) at least suggests this possibility. Like interventions that extend lifespan in *C. elegans*, the evolution of longer-lived mammals from shorter-lived ones often involves coupled reductions in both *α* and *β*, where mortality is both postponed and spread out over greater lengths of time. For instance, *α* at puberty decreases from 0.03 in laboratory mice to 0.0002 in humans, and subsequent mortality rate doubling time (an inverse measure of *β*) respectively increases from 0.27 to 8 years (Finch et al., 1990).

This raises the interesting question of whether evolution of greater longevity might also reduce *α* by extending healthspan and reduce *β* by ETL (fig. S6). In other words, the reduction of *β* in longer-lived mammals might only indirectly reflect reduced biological aging rate, which is rather directly reflected in the reduction of *α*. This coupled evolution of healthspan and gerospan expansion, and therefore *α* and *β*, could emerge from biological constraints present between mechanisms of development and aging (Gems and Kern, 2024) that exhibit evolutionary conservation. This would be consistent with the proportional scaling between life stages across mammalian species, such as that between ontogenetic span and adult lifespan (∼1:4) (Charnov, 1993). Additionally, the greater variability of gerospan than healthspan could arise from the late-life natural selection shadow, a key determinant of the evolution of aging (Medawar, 1952, Williams, 1957) that predicts greater optimization (thus, standardization) of early than later-life traits (fig. S6B).

In summary, these findings invert traditional conjectures about Gompertzian mortality: the rate parameter *β* reflects not biological aging rate but inter-individual variation in gerospan, the scale parameter *α* reflects less aging-independent mechanisms than the rate of aging itself, and both parameters reflect subpopulation-specific rather than population-wide traits. The approach used here, the combined analysis of individual and population aging, could prove similarly informative for understanding the biology of mortality patterns in higher organisms.

## Methods

### *C. elegans* culture and strains

*C. elegans* were maintained at 20°C using standard protocols (Brenner, 1974), on Nematode Growth Medium (NGM) plates seeded (2 days before use) with a bacterial food source (*Escherichia coli* OP50). 5-fluoro-2-deoxyuridine (FUDR), sometimes used to block progeny production, was not used in this study. Nematode strains used were: N2 (wild-type, hermaphrodite stock (Zhao et al., 2019)), GA1959 *daf-2(m577) III*, GA1960 *daf-2(e1368) III*, and GA1928 *daf-2(e1370) III*. All strains were raised from egg at 20°C on live *E. coli*, and transferred at L4 stage to the appropriate experimental conditions (15°C, 20°C or 25°C; with or without carbenicillin). Carbenicillin solution was added topically to plates one day before adding animals (further details below).

### Lifespan-only trials (Trials 1-2)

Nematodes were cultured throughout life in 60 mm Petri dishes (containing 10 mL of NGM) seeded with approximately 80 μL of *E. coli*, and where relevant, treated with 80 μL of 500 mM carbenicillin (Fisher Scientific Ltd, catalogue no. 12737149). At L4 stage (time 0 in all analyses), 30-40 animals were placed on each plate, with three plates per condition. Animals were transferred every two days during the reproductive period, and approximately every seven days thereafter. Scoring of survival was performed every two days. Animals showing no movement were gently touched with a platinum wire (worm pick) on the head and/or tail; those that showed no movement at all in response were scored as dead. Animals that died due to desiccation on the Petri dish wall, internal hatching of larvae, or rupture of internal tissues through the vulva, or that became contaminated by non-*E. coli* bacteria or fungi, or could not be found, were censored.

### Lifespan and healthspan trials (Trials 3-6)

Prior to the end of egg laying, nematodes were handled as for the lifespan-only trials, but with two plates containing 25 animals each, per condition. Following the end of egg laying, animals were transferred to individual wells of 24-well tissue culture plates, containing 2 mL of NGM and seeded with 3.5 μL of *E. coli* OP50, and where relevant, treated with 16 μL of 500 mM carbenicillin. Animals were subsequently transferred to fresh plates monthly, before media desiccation (plates were sealed with parafilm to delay desiccation, and to prevent bacterial/fungal contamination). Scoring of survival and censors were performed as above every 2-3 days, alongside scoring of locomotory class, and necropsy at death.

### Quantification of locomotory decline with age

Locomotory health class (belonging to H-span or G-span) was scored by classifying individuals into one of three classes, adapted from earlier systems (Hosono et al., 1980, Herndon et al., 2002): A – sinusoidal locomotion; B – non-sinusoidal locomotion; C – no locomotion. To accurately determine locomotory class, animals were gently touched on the tail with a platinum wire worm pick for up to 20 seconds to encourage movement, and additionally on the head as a final check. The duration spent in A class was defined as H-span, and the summed duration spent in B and C classes as G-span. Here, B and C classes were summed to improve data tractability, and to provide a definition of G-span that captures both early and late-stage functional declines.

### Necropsy analysis

Necropsy to define patterns of *E. coli*-associated pathology was performed by examining fresh corpses under a Leica MZ8 stereomicroscope (50x magnification). Scoring of swollen, bacterially-infected pharynxes (P), and uninfected, atrophied pharynxes (p) was performed as previously described (Zhao et al., 2017). Intestinal colonization (IC) with *E. coli* was scored where severe bacterial accumulation was observed in the anterior and/or posterior intestine. Such colonization presented as extreme lumenal distension by proliferating bacteria and/or colonization of the intestine beyond the lumenal barrier, with concomitant intestinal tissue degeneration and atrophy. Consistent across pharyngeal and intestinal tissues, sites of bacterial colonization exhibit a yellowish-brown color (as that of the *E. coli* lawn and colocalizing with RFP-labeled *E. coli*), translucent and uniform texture (loss of healthy tissue structures that otherwise appear dark, granular and opaque), and swollen/distended morphology (extensive proliferation of live *E. coli*). Images of representative examples of the three corpse subpopulations (P, pIC and pnIC) are presented in Fig. 4A and fig. S3B.

### Microscopy

Microscopy slides were prepared by placing individual nematode corpses in a small drop of M9 buffer on 2% agar pads, under glass coverslips. Brightfield images were captured using an ApoTome.2 Zeiss microscope with a Hamamatsu digital camera C13440 ORCA-Flash4.0 V3 and Zen software, at 100x total magnification, with 125 ms exposure time and 1.1 V illumination intensity. The presence of *E. coli* OP50-RFP in the pharynx was assayed using the mRF12 channel (excitation: 577–604□nm; emission: 612□nm) at the same magnification, with 750 ms exposure time and 75% LED intensity. Brightness and contrast were adjusted equally across the entire image, and where applicable applied equally to controls. Brightfield and RFP epifluorescence necropsy images were overlaid in ImageJ and backgrounds removed with Adobe Express (online tool). The maximum intensity threshold of RFP channel images was adjusted in ImageJ from 255 to 70 for all images.

### Mortality deconvolution (subpopulation) analyses

Age-specific survival proportions for full (not deconvolved into subpopulations) populations were obtained by conventional Kaplan-Meier analysis (including censors), from which age-specific mortality pseudofrequencies (which sum to 1) were calculated. For each age, these pseudofrequencies were partitioned into subpopulation pseudofrequencies, weighted by the proportion of total mortality at that age belonging to each subpopulation. Standard survival and mortality analyses of subpopulations were then performed utilizing these pseudofrequencies (in place of conventional mortality frequency), to enable unbiased inclusion of censor data in subpopulation analyses. Survival analyses in Fig. 4B-D were performed in this manner. Contributions of subpopulation changes to the Gompertz parameters in Fig. 5A-D were similarly performed on simulated (mortality pseudofrequency-derived) survival data. Simulated survival data was generated for each component change (in subpopulation prevalence or lifespan), by accordingly combining simulated control and treatment subpopulations. Specifically, changes in subpopulation prevalence were simulated by combining treatment cohort subpopulation prevalence with control subpopulation lifespans (based on principles of parsimony, Fig. 4E-F), and changes in lifespan were simulated by combining treatment cohort subpopulation lifespan with control subpopulation prevalence, for each subpopulation at a time. The sum of these component changes closely predicts the true change between control and treatment cohorts (fig. S5A). Contributions of component changes to *α* and *β* were then estimated as the change in, respectively, lifespan of the 10% shortest-lived individuals and lifespan variance, which strongly predict *α* and *β* across these 12 non-antibiotic cohorts (fig. S5B-C) and whose component sum of these changes predicts their true change between control and treatment cohorts (fig. S5D-E).

### Statistics, software and data handling

Statistical tests were performed using JMP Pro (SAS Institute, Inc.), except for Gompertz parameter estimation and assessment of statistical differences between them, which were performed using WinModest (Pletcher, 1999). Right censors were included in all Kaplan-Meier and WinModest analyses. Specific statistical tests and associated methodological details are described in the respective figure/table captions. Notation of statistical significance is as follows: *p* > 0.05, * *p* ≤ 0.05, ** *p* ≤ 0.01, *** *p* ≤ 0.001, **** *p* ≤ 0.0001. Necropsy image capture, processing and editing was performed using Zen software and ImageJ.

Analyses of treatment effects on lifespan (fig. S1, table S1) were performed on the pool of all 6 trials, while analyses relating to locomotory healthspan and gerospan (Fig. 1-3, fig. S2, table S2-4) were performed on the pool of 4 trials (Trials 3-6; see Methods above), in which locomotory senescence was quantified. Subpopulation analyses utilizing necropsy data (Fig. 4D-I, 5A-D, fig. S3A, S4-5, table S5-6) were similarly performed on this pool of 4 trials (or 3 trials, Trials 4-6, for N2 cohorts), in which necropsy was performed. Further information and summary statistics for individual trials are provided in table S1.

## Data availability

Raw data for all 6 trials is provided, including lifespan, locomotory healthspan and gerospan, and necropsy subpopulation type for all individuals where measured (*n*=8,830).

## Supplementary Material

Supplementary Figures, Supplementary Tables.

## Acknowledgments

We thank A. Vere-Hopegood, A. Zhang and H. Chapman for minor research contributions, J.P. de Magalhães, S.D. Pletcher, J. Labbadia and T. Niccoli for useful discussion, and Y. Zhao and S.D. Pletcher for comments on the manuscript. Some strains were provided by the Caenorhabditis Genetics Center, which is funded by the NIH Office of Research Infrastructure Programs (P40 OD010440).

## Author contributions

D.G. supervised the project. B.Z. and D.G. conceived the project, designed the experiments and data analysis, and wrote the manuscript. B.Z. performed the experiments and analyzed the data.

## References

Bansal, A., Zhu, L. J., Yen, K. & Tissenbaum, H. A. 2015. Uncoupling lifespan and healthspan in Caenorhabditis elegans longevity mutants. Proc Natl Acad Sci U S A, 112, E277–86.

Blagosklonny, M. V. 2010. Why men age faster but reproduce longer than women: mTOR and evolutionary perspectives. Aging (Albany NY), 2, 265–73.

Brenner, S. 1974. The genetics of Caenorhabditis elegans. Genetics, 77, 71–94.

Charnov, E. L. 1993. Life History Invariants. Some Explorations of Symmetry in Evolutionary Ecology, Oxford, Oxford University Press.

Churgin, M. A., Jung, S. K., Yu, C. C., Chen, X., Raizen, D. M. & Fang-YEN, C. 2017. Longitudinal imaging of Caenorhabditis elegans in a microfabricated device reveals variation in behavioral decline during aging. Elife, 6.

Driver, C. 2001. The Gompertz function does not measure ageing. Biogerontology, 2, 61–5.

Driver, C. 2003. A further comment on why the Gompertz plot does not measure aging. Biogerontology, 4, 325–7.

Eakin, T., Shouman, R., Qi, Y., Liu, G. & Witten, M. 1995. Estimating parametric survival model parameters in gerontological aging studies: methodological problems and insights. J Gerontol A Biol Sci Med Sci, 50, B166–76.

Finch, C. E., Pike, M. C. & Witten, M. 1990. Slow mortality rate accelerations during aging in some animals approximate that of humans. Science, 249, 902–905.

Garigan, D., Hsu, A. L., Fraser, A. G., Kamath, R. S., Ahringer, J. & Kenyon, C. 2002. Genetic analysis of tissue aging in Caenorhabditis elegans: a role for heat-shock factor and bacterial proliferation. Genetics, 161, 1101–1112.

Gems, D. & Kern, C. C. 2024. Biological constraint, evolutionary spandrels and antagonistic pleiotropy. Ageing Res Rev, 101, 102527.

Gems, D., Sutton, A. J., Sundermeyer, M. L., Larsen, P. L., Albert, P. S., King, K. V., Edgley, M. & Riddle, D. L. 1998. Two pleiotropic classes of daf-2 mutation affect larval arrest, adult behavior, reproduction and longevity in Caenorhabditis elegans. Genetics, 150, 129–155.

Gompertz, B. 1825. Of the nature of the function expressive of the law of human mortality, and on a new mode of determining the value of Life Contingencies. Philos Trans R Soc London Ser A, 57, 513–585.

Greenwood 1928. “Laws” of Mortality from the Biological point of view. J Hyg (Lond), 28, 267–94.

Guo, J., Huang, X., Dou, L., Yan, M., Shen, T., Tang, W. & Li, J. 2022. Aging and aging-related diseases: from molecular mechanisms to interventions and treatments. Signal Transduct Target Ther, 7, 391.

Hahm, J., Kim, S., Diloreto, R., Shi, C., Lee, S., Murphy, C. & Nam, H. 2015. C. elegans maximum velocity correlates with healthspan and is maintained in worms with an insulin receptor mutation. Nat Commun, 6, 8919.

Herndon, L. A., Schmeissner, P. J., Dudaronek, J. M., Brown, P. A., Listner, K. M., Sakano, Y., Paupard, M. C., Hall, D. H. & Driscoll, M. 2002. Stochastic and genetic factors influence tissue-specific decline in ageing C. elegans. Nature, 419, 808–814.

Hosono, R., Sato, Y., Aizawa, S. I. & Mitsui, Y. 1980. Age-dependent changes in mobility and separation of the nematode Caenorhabditis elegans. Exp Gerontol, 15, 285–9.

Huang, C., Xiong, C. & Kornfeld, K. 2004. Measurements of age-related changes of physiological processes that predict lifespan of Caenorhabditis elegans. Proc Natl Acad Sci U S A., 101, 8084–8089.

Hughes, B. G. & Hekimi, S. 2016. Different mechanisms of longevity in long-lived mouse and Caenorhabditis elegans mutants revealed by statistical analysis of mortality rates. Genetics, 204, 905–920.

Jones, O. R., Scheuerlein, A., Salguero-Gómez, R., Camarda, C. G., Schaible, R., Casper, B. B., Dahlgren, J. P., Ehrlén, J., García, M. B., Menges, E. S., Quintana-Ascencio, P. F., Caswell, H., Baudisch, A. & Vaupel, J. W. 2014. Diversity of ageing across the tree of life. Nature, 505, 169–73.

Kenyon, C., Chang, J., Gensch, E., Rudener, A. & Tabtiang, R. 1993. A C. elegans mutant that lives twice as long as wild type. Nature, 366, 461–464.

Kirkwood, T. B., Feder, M., Finch, C. E., Franceschi, C., Globerson, A., Klingenberg, C. P., Lamarco, K., Omholt, S. & Westendorp, R. G. 2005. What accounts for the wide variation in life span of genetically identical organisms reared in a constant environment? Mech Ageing Dev, 126, 439–43.

Klass, M. R. 1977. Aging in the nematode Caenorhabditis elegans: major biological and environmental factors influencing life span. Mech Ageing Develop, 6, 413–429.

Koopman, J. J., Rozing, M. P., Kramer, A., De JAGER, D. J., Ansell, D., De MEESTER, J. M., Prütz, K. G., Finne, P., Heaf, J. G., Palsson, R., Kramar, R., Jager, K. J., Dekker, F. W. & Westendorp, R. G. 2011. Senescence rates in patients with end-stage renal disease: a critical appraisal of the Gompertz model. Aging Cell, 10, 233–8.

Masoro, E. J. 2006. Caloric restriction and aging: controversial issues. J Gerontol A Biol Sci Med Sci, 61, 14–9.

Medawar, P. B. 1952. An Unsolved Problem Of Biology, London, H.K. Lewis.

Mueller, L. D., Nusbaum, T. J. & Rose, M. R. 1995. The Gompertz equation as a predictive tool in demography. Exp Gerontol, 30, 553–69.

Newell Stamper, B. L., Cypser, J. R., Kechris, K., Kitzenberg, D. A., Tedesco, P. M. & Johnson, T. E. 2018. Movement decline across lifespan of Caenorhabditis elegans mutants in the insulin/insulin-like signaling pathway. Aging Cell, 17, e12704.

Pletcher, S. 1999. Model fitting and hypothesis testing for age-specific mortality data. J Evol Bio., 12, 430–439.

Podshivalova, K., Kerr, R. & Kenyon, C. 2017. How a mutation that slows aging can also disproportionately extend end-of-life decrepitude. Cell Reports, 19, 441–450.

Rozing, M. P. & Westendorp, R. G. 2008. Parallel lines: nothing has changed? Aging Cell, 7, 924–7.

Sacher, G. A. 1977. Life table modification and life prolongation. In: Finch, C. E. & Hayflick, L. (eds.) Handbook of the Biology of Aging. New York: Van Norstrand Reinhold.

Shouman, R. & Witten, M. 1995. Survival estimates and sample size: what can we conclude? J Gerontol A Biol Sci Med Sci, 50, B177–85.

Statzer, C., Reichert, P., Dual, J. & Ewald, C. Y. 2022. Longevity interventions temporally scale healthspan in Caenorhabditis elegans. iScience, 25, 103983.

Stroustrup, N., Anthony, W. E., Nash, Z. M., Gowda, V., Gomez, A., López-Moyado, I. F., Apfeld, J. & Fontana, W. 2016. The temporal scaling of Caenorhabditis elegans ageing. Nature, 530, 103–107.

Uno, M., Nono, M., Takahashi, C., Kishimoto, S., Okabe, E., Yamamoto, T. & Nishida, E. 2025. A transition from interindividual uniformity to diversity in appearance and transcriptional features at midlife in Caenorhabditis elegans. Genes Cells, 30, e13187.

Vaupel, J. W., Carey, J. R., Christensen, K., Johnson, T. E., Yashin, A. I., Holm, N. V., Iachine, I. A., Kannisto, V., Khazaeli, A. A., Liedo, P., Longo, V. D., Zeng, Y., Manton, K. G. & Curtsinger, J. W. 1998. Biodemographic trajectories of longevity. Science, 280, 855–860.

Vaupel, J. W., Manton, K. G. & Stallard, E. 1979. The impact of heterogeneity in individual frailty on the dynamics of mortality. Demography, 16, 439–54.

Williams, G. C. 1957. Pleiotropy, natural selection and the evolution of senescence. Evolution, 11, 398–411.

Yashin, A. I., Ukraintseva, S. V., Boiko, S. I. & Arbeev, K. G. 2002. Individual aging and mortality rate: how are they related? Soc Biol, 49, 206–17.

Zhang, W., Sinha, D., Pittman, W., Hvatum, E., Stroustrup, N. & Pincus, Z. 2016. Extended twilight among isogenic C. elegans causes a disproportionate scaling between lifespan and health. Cell Syst, 3, 333–345.

Zhao, Y., Gilliat, A. F., Ziehm, M., Turmaine, M., Wang, H., Ezcurra, M., Yang, C., Phillips, G., Mcbay, D., Zhang, W. B., Partridge, L., Pincus, Z. & Gems, D. 2017. Two forms of death in aging Caenorhabditis elegans. Nat Commun, 8, 15458.

Zhao, Y., Wang, H., Poole, R. J. & Gems, D. 2019. A fln-2 mutation affects lethal pathology and lifespan in C. elegans. Nat Commun, 10, 5087.

Zhao, Y., Zhang, B., Marcu, I., Athar, F., Wang, H., Galimov, E. R., Chapman, H. & Gems, D. 2021. Mutation of daf-2 extends lifespan via tissue-specific effectors that suppress distinct life-limiting pathologies. Aging Cell, 20, e13324.

