## Supplemental Data for "Slow Gompertzian aging in long-lived *C. elegans* results from expansion of decrepitude, not decelerated aging"

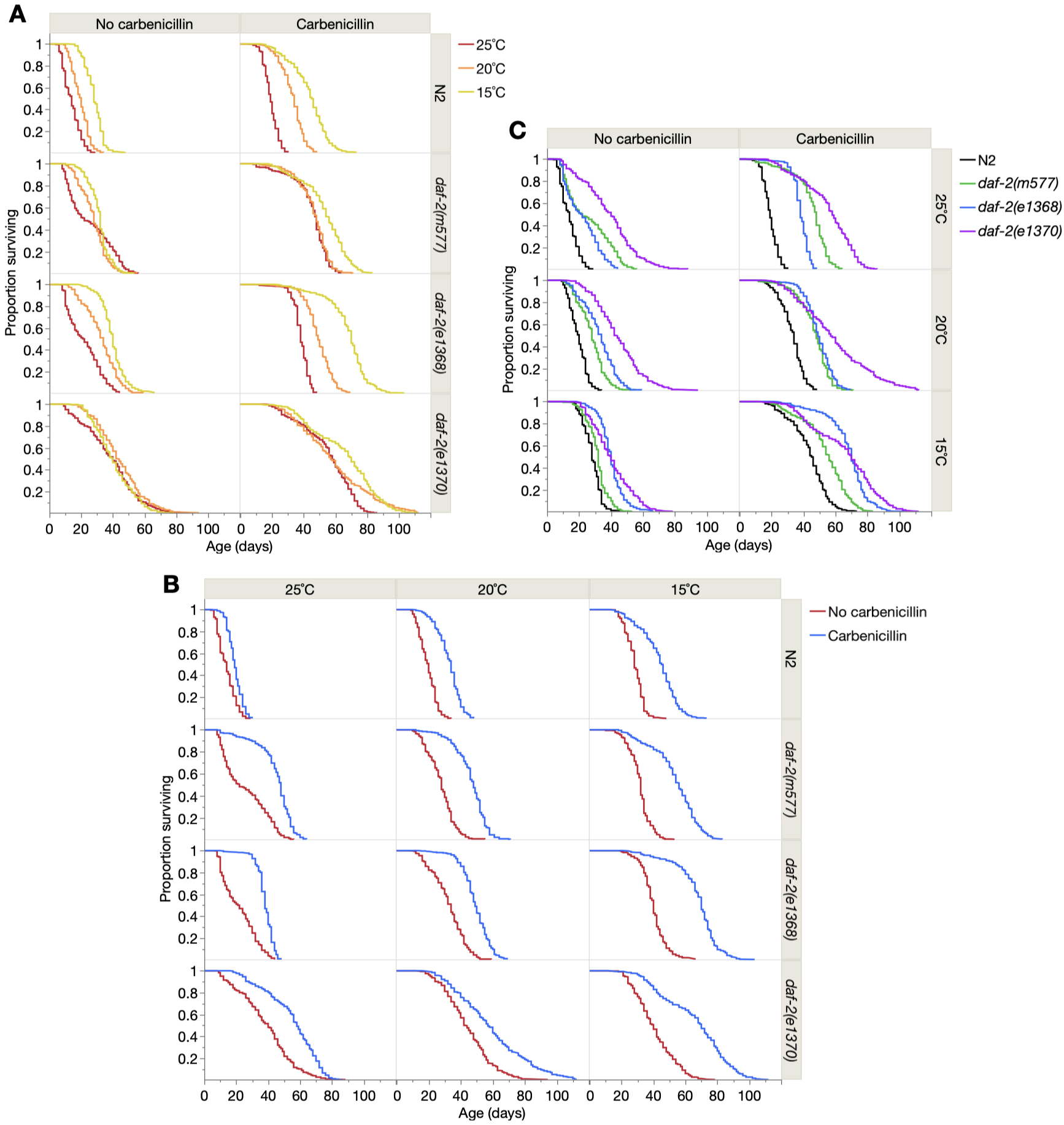

Fig. S1.

**Kaplan-Meier survival curves for all cohorts**. Effects of (**A**) reducing temperature, (**B**) antibiotic treatment, and (**C**) *daf-2(rf)*. N2, wild-type. Pool of all 6 trials. Statistical comparisons (*p* values, log rank test) are presented in table S1. A previous study found that culture of *daf-2(e1370)* on a different antibiotic (gentamycin) did not increase lifespan, and even slightly shortened it, suggesting that this mutation might confer full resistance to life-shortening effects of live *E. coli* (*32*). Our findings using carbenicillin (B) imply that all 3 *daf-2* mutants succumb to effects of proliferating *E. coli* in later life, likely reflecting inevitable, eventual immunosenescence.

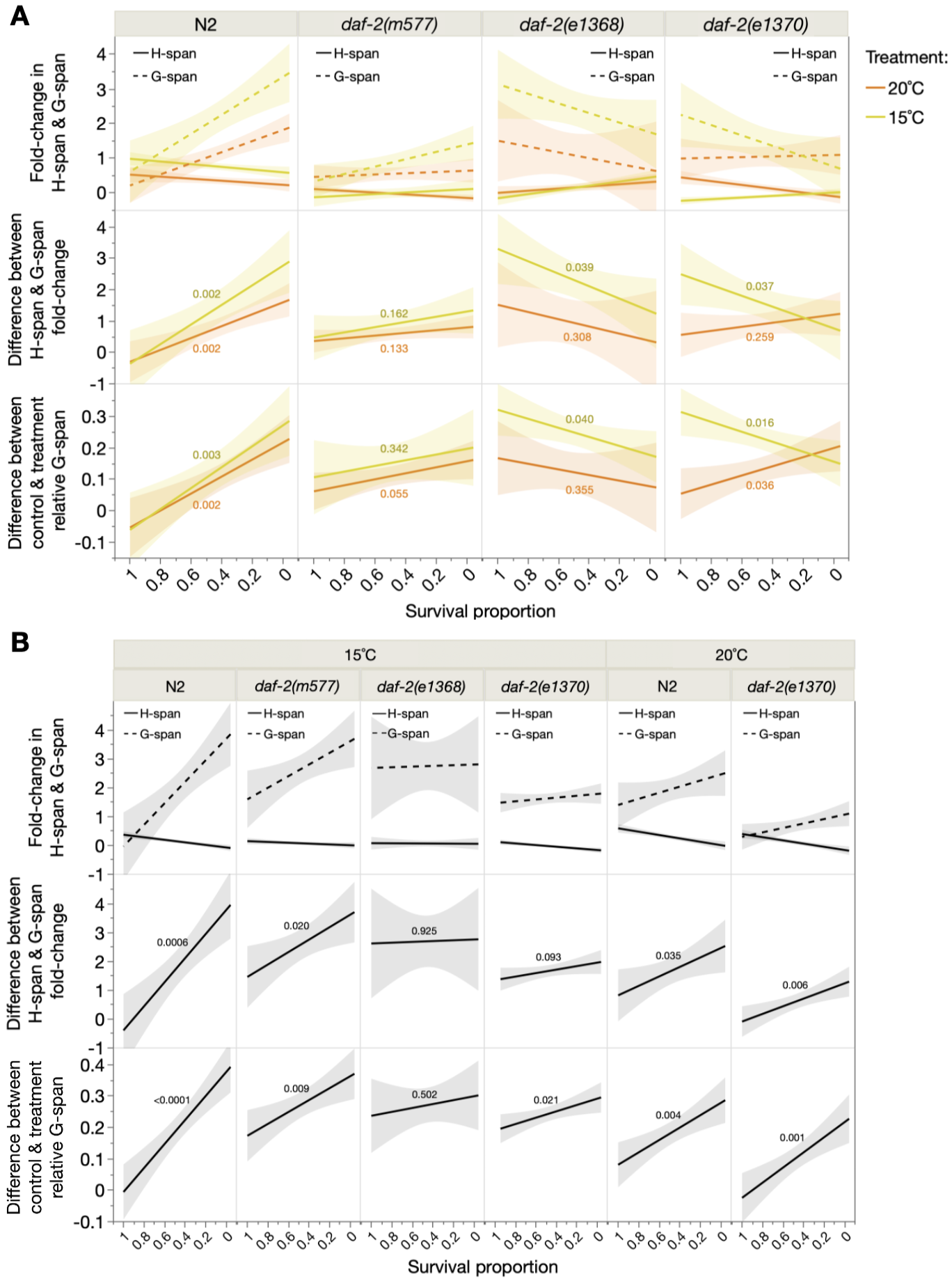

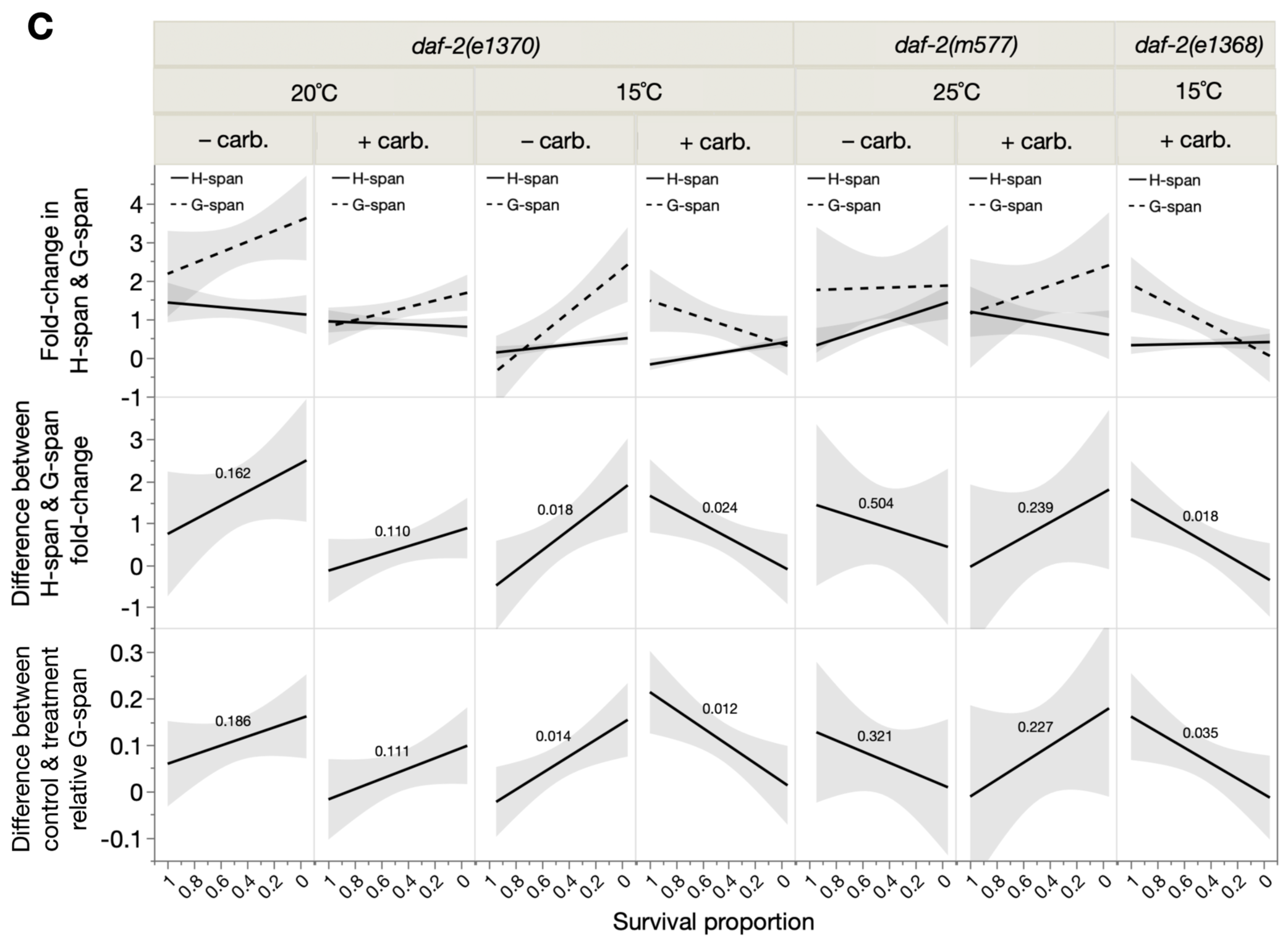

Fig. S2.

**Extended twilight longevity (ETL) increases gerospan in longer-lived individuals**. All longevity treatments causing simultaneous *β* decrease and G-span^rel^ increase are presented, from three treatment categories: (**A**) low temperature, (**B**) carbenicillin and (**C**) *daf-2(rf)*. N2: wild-type. The **top row of panels** plots the fold-change of absolute H-span and G-span (in response to treatment) over survival proportion (i.e. x-axis left: shorter-lived individuals, x-axis right: longer-lived individuals); in ETL, G-span fold-change is greater than H-span fold-change at lower survival proportions (longer-lived individuals). This difference between G-span and H-span fold-change across survival proportions is plotted and evaluated in the **middle row of panels**; in ETL, this yields a positive relationship, whereas inverted ETL yields a negative relationship. The **bottom row of panels** plots and evaluates the difference between control and treatment G-span^rel^ (from Fig. 2B-G, lower panels) over survival proportion; in ETL, this yields a positive relationship, whereas inverted ETL yields a negative relationship. All panels show a least-squares linear regression with shaded 95% confidence regions, and associated F-test *p*-values overlaid for middle and bottom panels. We note these regressions may be limited in power by small sample size (8 ≤ *n* ≤ 20), due to binning and averaging of individual survival proportions (0.05 survival proportion bins) and selective inclusion of only those bins represented in both control and treatment cohorts. However, regardless of statistical significance, all relationship directions (positive or negative) agree with those in Fig. 2B-G, which perform regressions using all (unbinned) individuals (104 ≤ *n* ≤ 173).

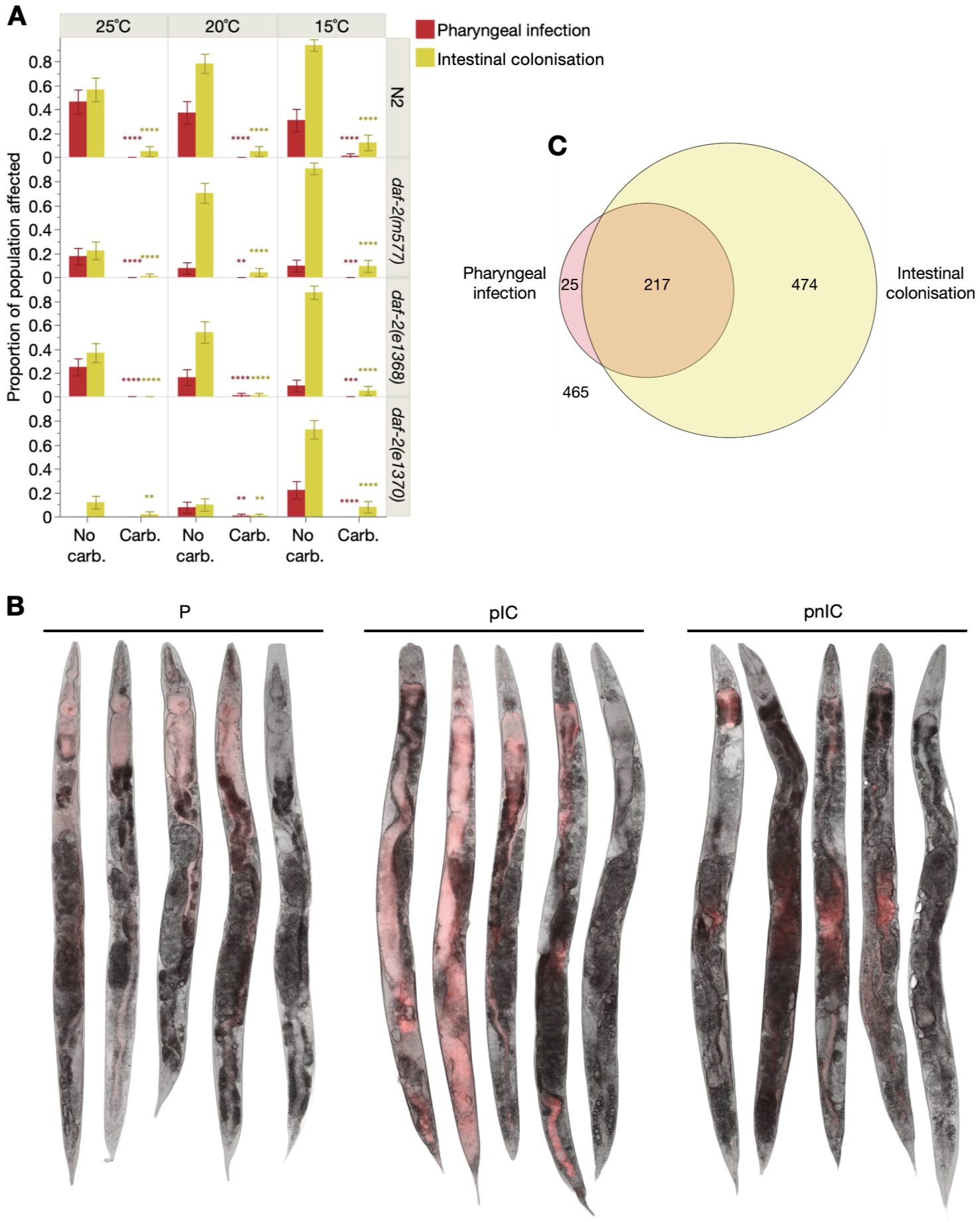
Fig. S3.

**Pharyngeal and intestinal tissues are colonized by dietary *E. coli* during aging**. (**A**) Antibiotic (carbenicillin) treatment abolishes pharyngeal and intestinal colonization, as scored visually under a Leica MZ8 stereomicroscope (50x total magnification), based on tissue color, texture and morphology, as described in the methods and performed previously (*20*). 95% confidence intervals are shown. Differences in proportion of the population affected were assessed by Pearson’s chi-squared test. Statistical significance notation: ns *p* > 0.05, * *p* ≤ 0.05, ** *p* ≤ 0.01, *** *p* ≤ 0.001, **** *p* ≤ 0.0001. (**B**) Representative images of the range of corpse presentations for each subpopulation (P, pIC, pnIC), taken from a single population (*n*=45) of nematodes fed throughout life and colonized by RFP-expressing *E. coli* (first four examples in each subpopulation). The fifth (last) example in each subpopulation is of individuals fed standard *E. coli* (lacking RFP expression) throughout life, showing that RFP signal is attributable to bacteria rather than nematode autofluorescence, and that presentation of the described pathologies does not differ between standard and RFP-expressing *E. coli* diets. From left to right, lifespan in days for these specific individuals: P (12, 14, 16, 20, 16), pIC (12, 14, 20, 22, 26), pnIC (16, 20, 24, 34, 30). See Methods for detailed microscopy methodology. Interestingly, epifluorescence microscopy of RFP-expressing *E. coli* reveals its uterine and/or tumoral colonization in pnIC individuals, which is less readily visible under a dissecting stereomicroscope than pharyngeal and intestinal colonization. (**C**) Venn diagram showing that most P individuals simultaneously have intestinal colonization by bacteria.

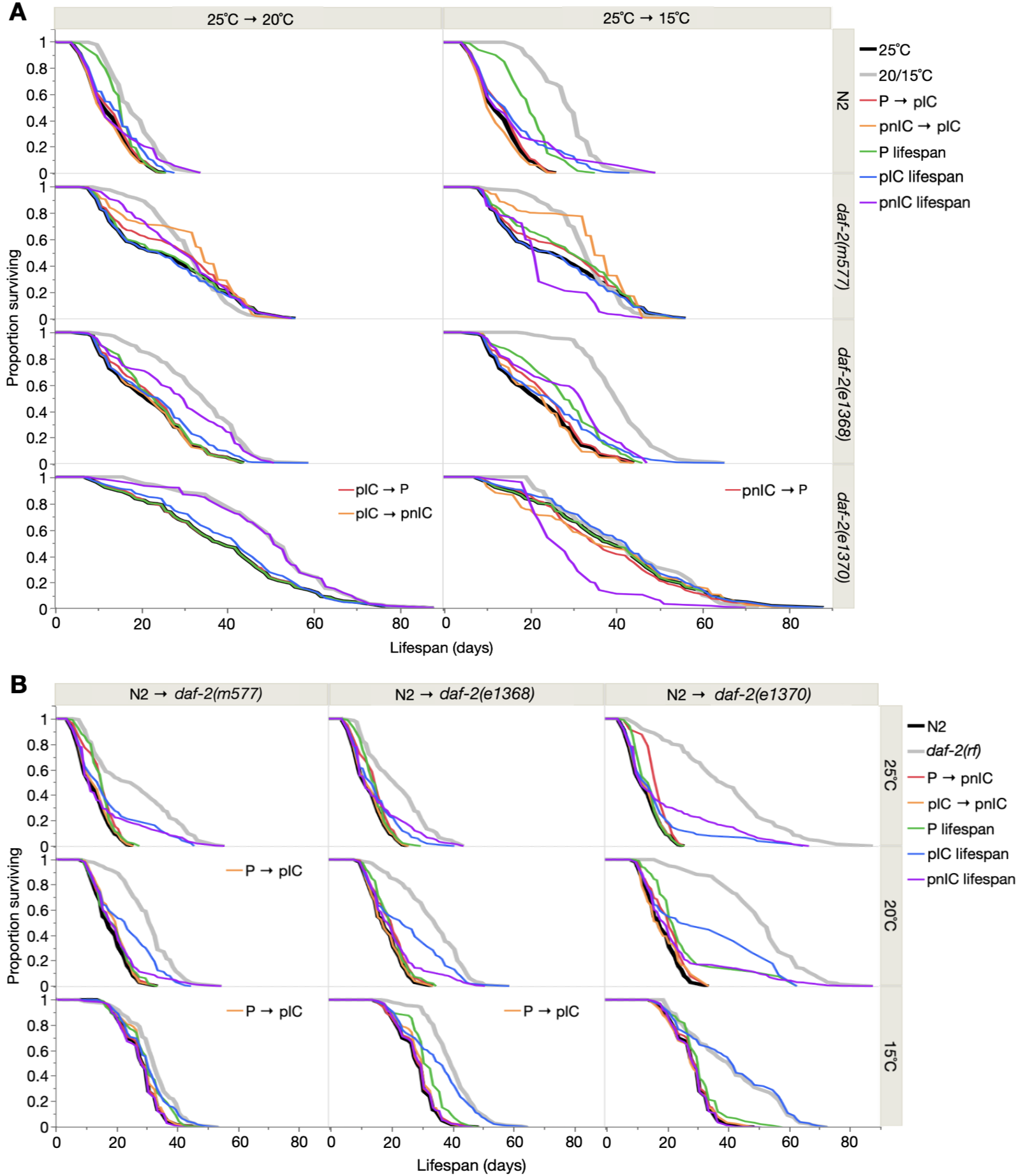

**Fig. S4.**

**Effect of life-extending treatments on subpopulation prevalence and lifespan.** Effect of (**A**) low temperature and (**B**) *daf-2(rf)* treatments. Black and grey cohorts are control and treatment cohorts, respectively. Red and orange cohorts are the component effects of changing control subpopulation prevalence to that of the treatment cohort, based on principles of parsimony (see Figure 4E-F), while retaining control subpopulation lifespans. Green, blue and purple cohorts are the effects of changing control subpopulation lifespan to that of the treatment cohort, while retaining control subpopulation prevalence, for P, pIC and pnIC, respectively. Survival proportions were obtained from Kaplan-Meier survival analysis using mortality pseudofrequencies (in place of frequency) to account for censors (see Methods). Sub-legends reflect different subpopulation prevalence changes for those specific treatments, similarly based on principles of parsimony.

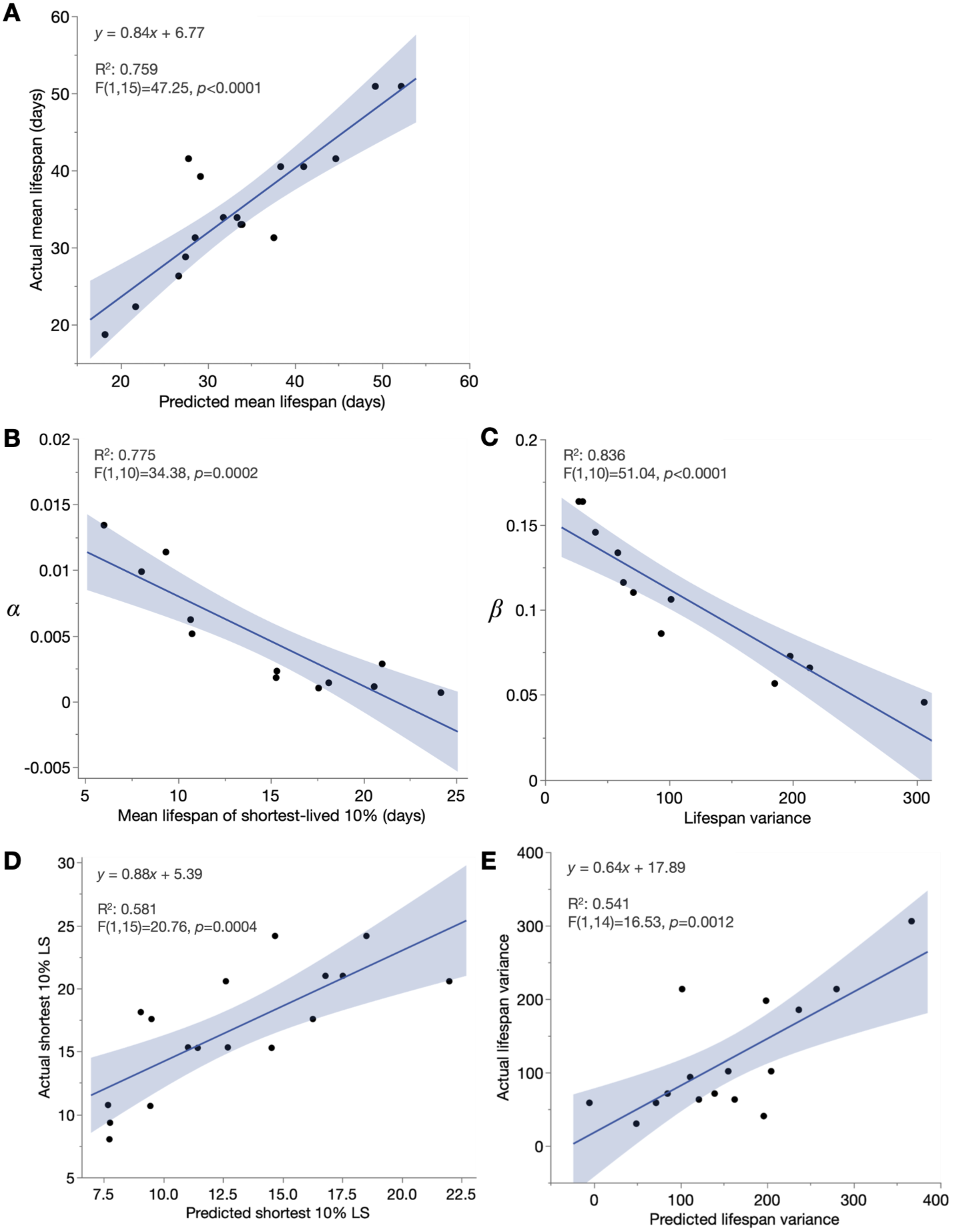

**Fig. S5.**

**Methodological background of determining relative contributions to the Gompertz parameters of subpopulation changes.** (**A**) Mean lifespan in the 17 treatments (low temperature and *daf-2(rf)*; non-antibiotic background) is strongly predicted by the sum of simulated component changes (in subpopulation prevalence and lifespan) (see Methods for analysis details). (**B**-**C**) Mean lifespan of 10% shortest-lived individuals and lifespan variance are strong respective predictors of *α* and *β* in the 12 non-antibiotic cohorts under analysis. (**D**-**E**) Simulated component changes (in subpopulation prevalence and lifespan) predict true changes in mean lifespan of the 10% shortest-lived individuals, and lifespan variance, in the 17 treatments (low temperature and *daf-2(rf)*; non-antibiotic background). In (E) one outlier treatment (*daf-2(e1370)* at 20˚C) was excluded, with datapoints x=754, y=198 (Robust Fit Outliers analysis in JMP Pro, Huber method, *K*=4).

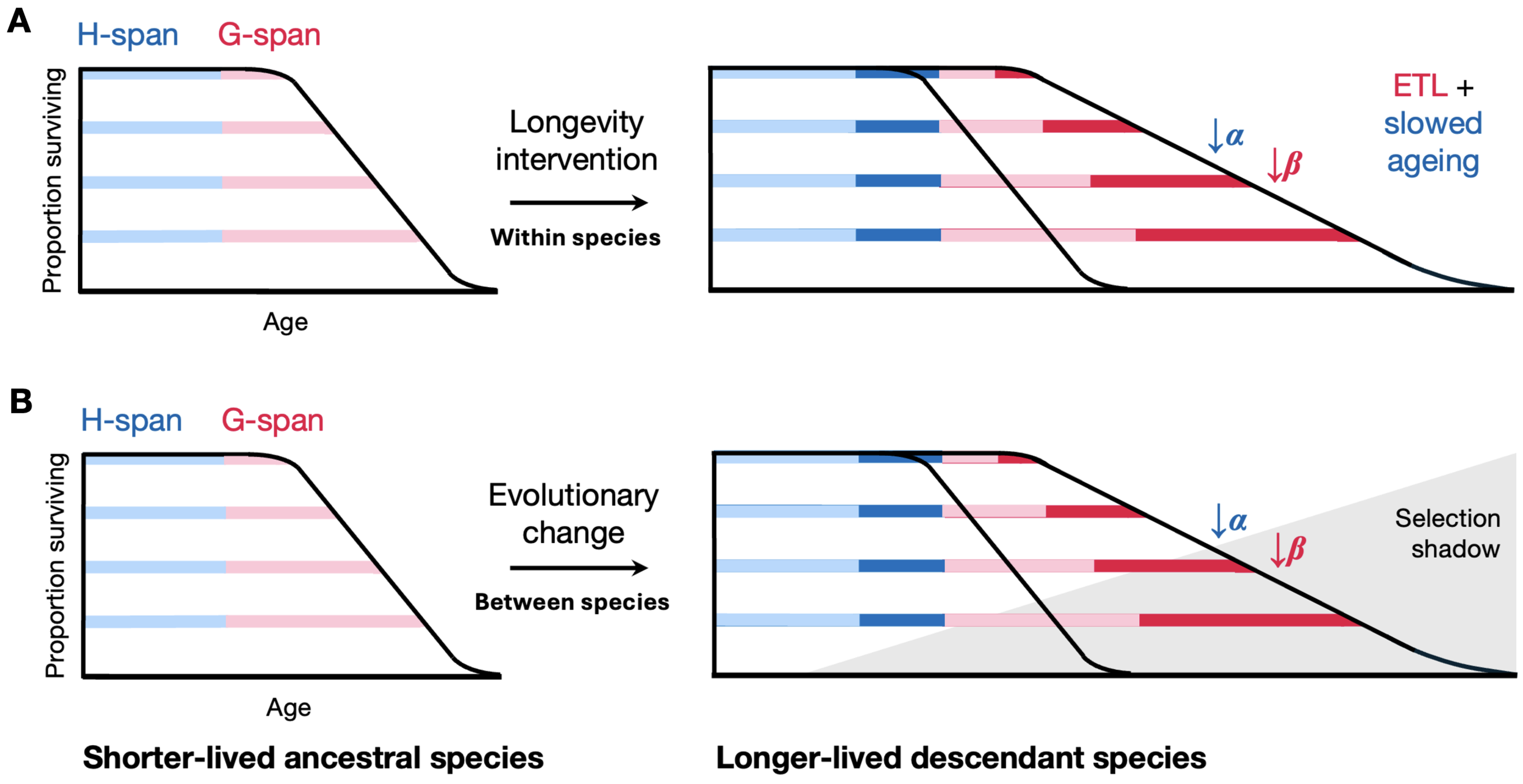

Fig. S6.
Potential role for ETL and healthspan extension in the evolution of longevity. Schematic of the biological basis of the Gompertz parameters within and hypothetically, between, species. (A) Reduction of *α* by healthspan (H-span) extension and reduction of *β* by extended twilight longevity (ETL). Each panel depicts variation in health and lifespan in 4 representative individuals; end of bar: point of death, as bordered by the survival curve. Here, as demonstrated in *C. elegans*, the increased variance in lifespan that reduces *β* arises from inter-individually variable expansion of gerospan (G-span), as part of the phenotypic variability that emerges and progressively increases during aging. This simplified representation shows no inter-individual variation or change in healthspan (H-span); more accurately, H-span also varies between individuals and increases with lifespan, but to a lesser magnitude than G-span (*21*). (B) Reduction in *α* and *β* during evolution of increased longevity (hypothetical scheme). Here again, reduced *β* reflects increased lifespan variance due to inter-individually variable expansion of G-span, arising from the selection shadow (reduced force of selection) in later life on the evolved genome. This evolutionary ETL is coupled to a less variable expansion of H-span, corresponding to slowed aging, that reduces *α*. For simplicity, again no inter-individual variation in H-span is depicted. In neither (A) nor (B) is *β* a direct metric of biological aging rate; we suggest it is instead *α* that is.

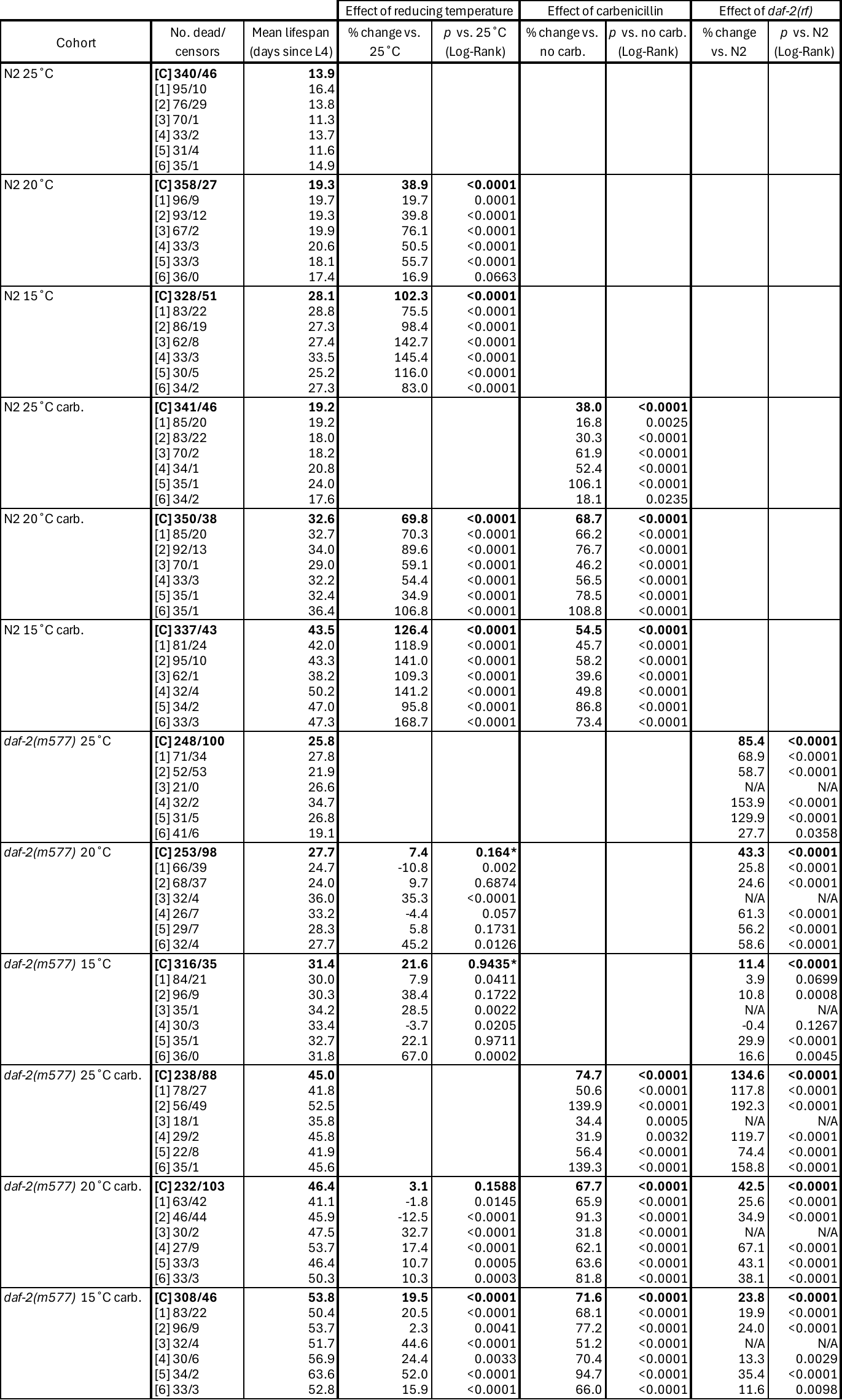

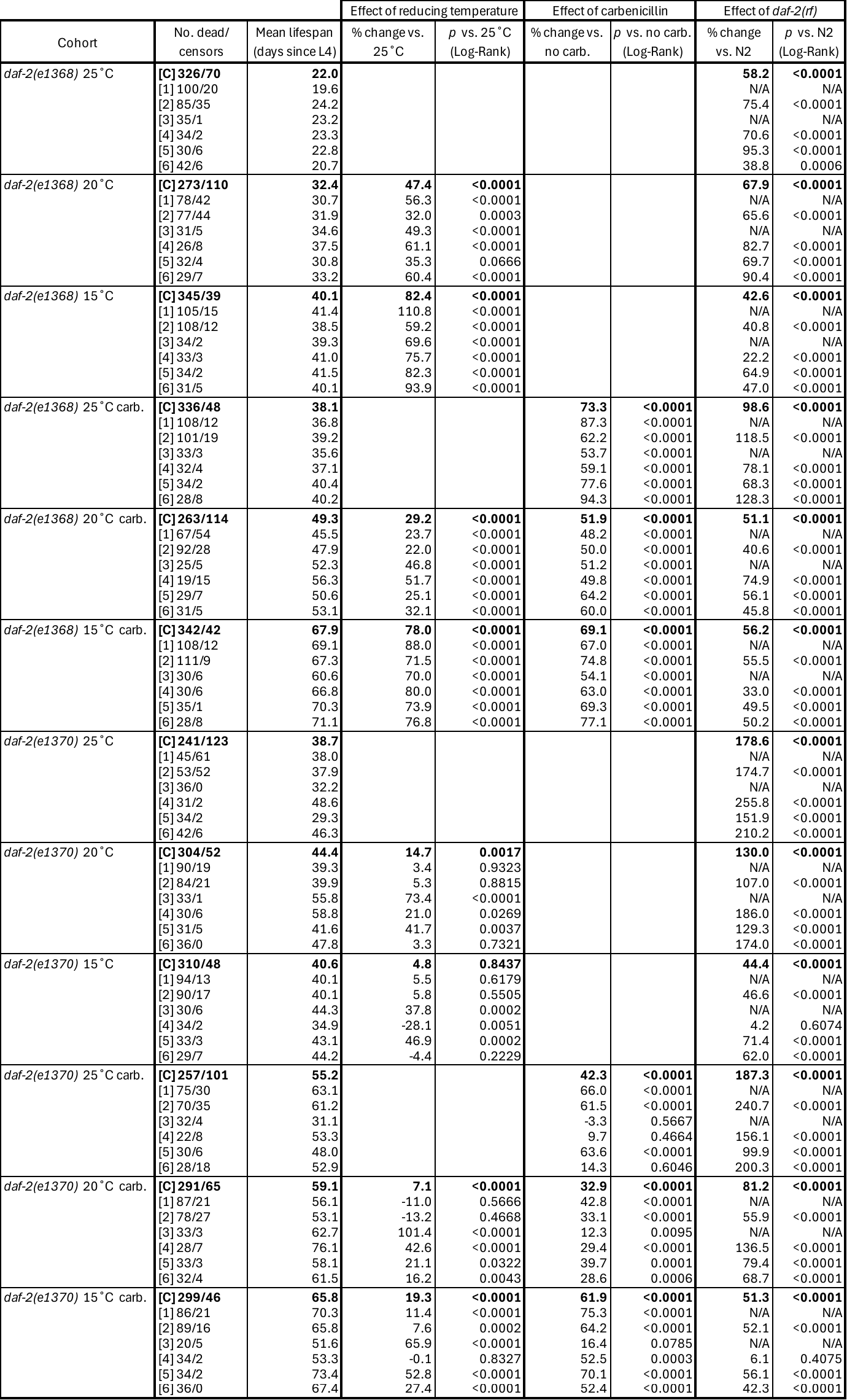

Table S1.

**Lifespan data for all cohorts and trials, and effect on lifespan of reducing temperature, antibiotic treatment and *daf-2(rf)*.** N2, wild-type. [C], combined (pooled) data from all trials; [*n*], trial number; carb., carbenicillin. *, cohorts where Log-Rank *p*>0.05, due to crossing survival curves and greater changes in early than late mortality; the Wilcoxon test is more appropriate to detect lifespan differences in these conditions (*p*<0.0015; data not shown). The 6 trials were performed sequentially over 34 months. In trials 1-3, experiments for the 24 cohorts were not always performed together due to practical constraints. In this table, intra-trial comparisons between such cohorts were therefore withheld, and these exclusions (affecting a small proportion of within-trial comparisons: 30/276) are labelled (N/A). However, all analyses performed in this study utilize pooled data from multiple trials (see Methods); in these analyses, the above-excluded comparisons were included, given that pooled analyses by nature compare different trials, and additionally dilute inter-trial noise.

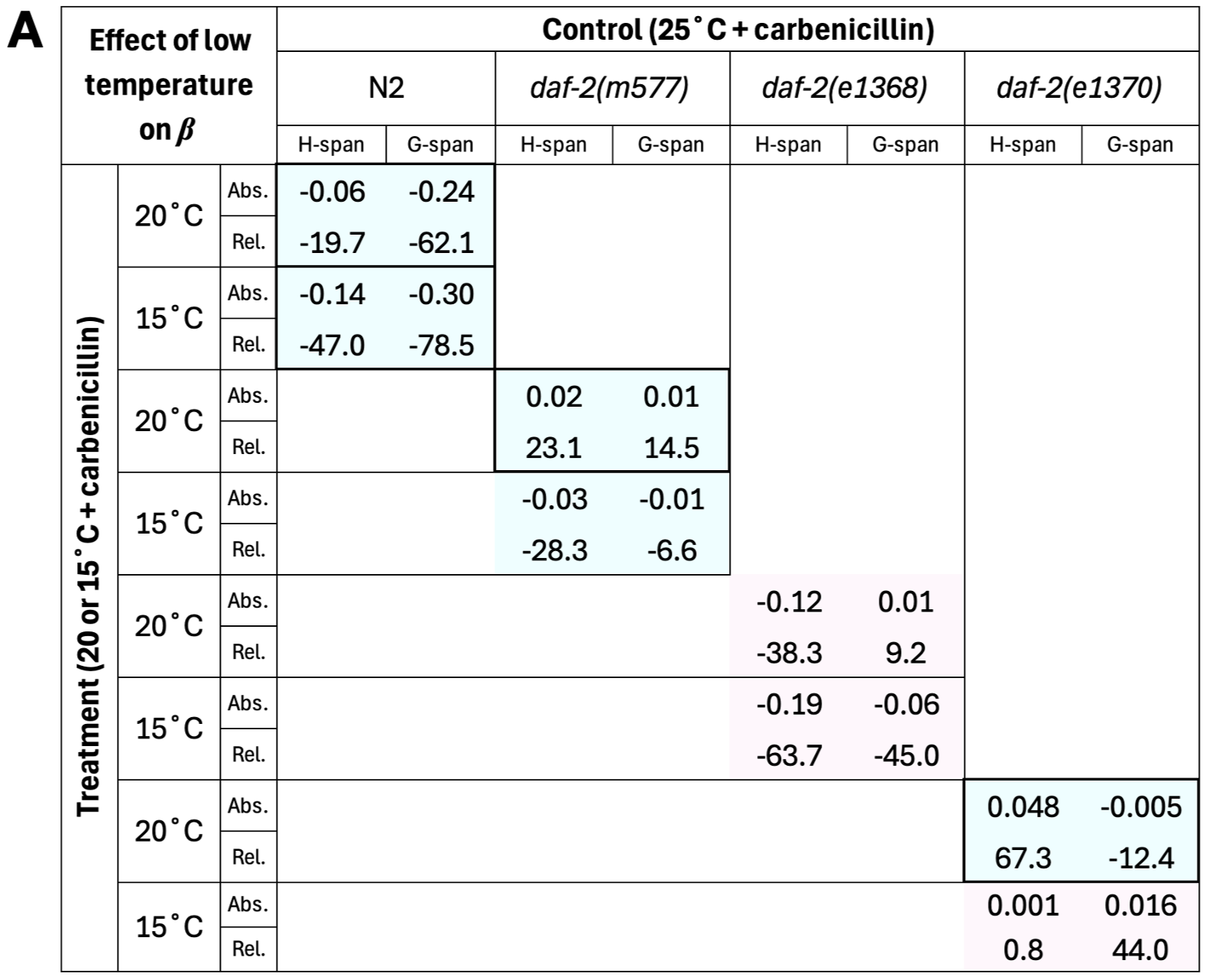

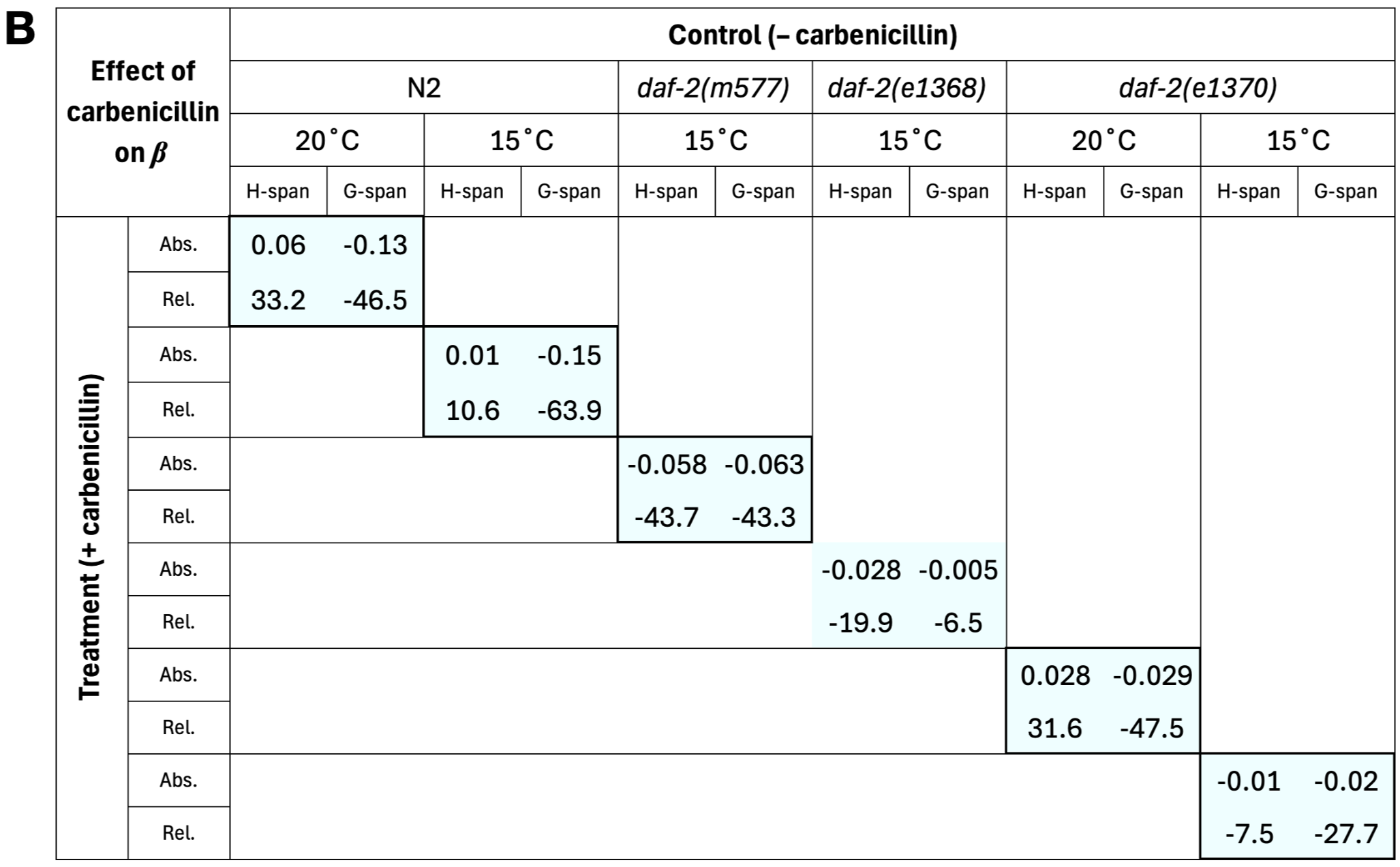

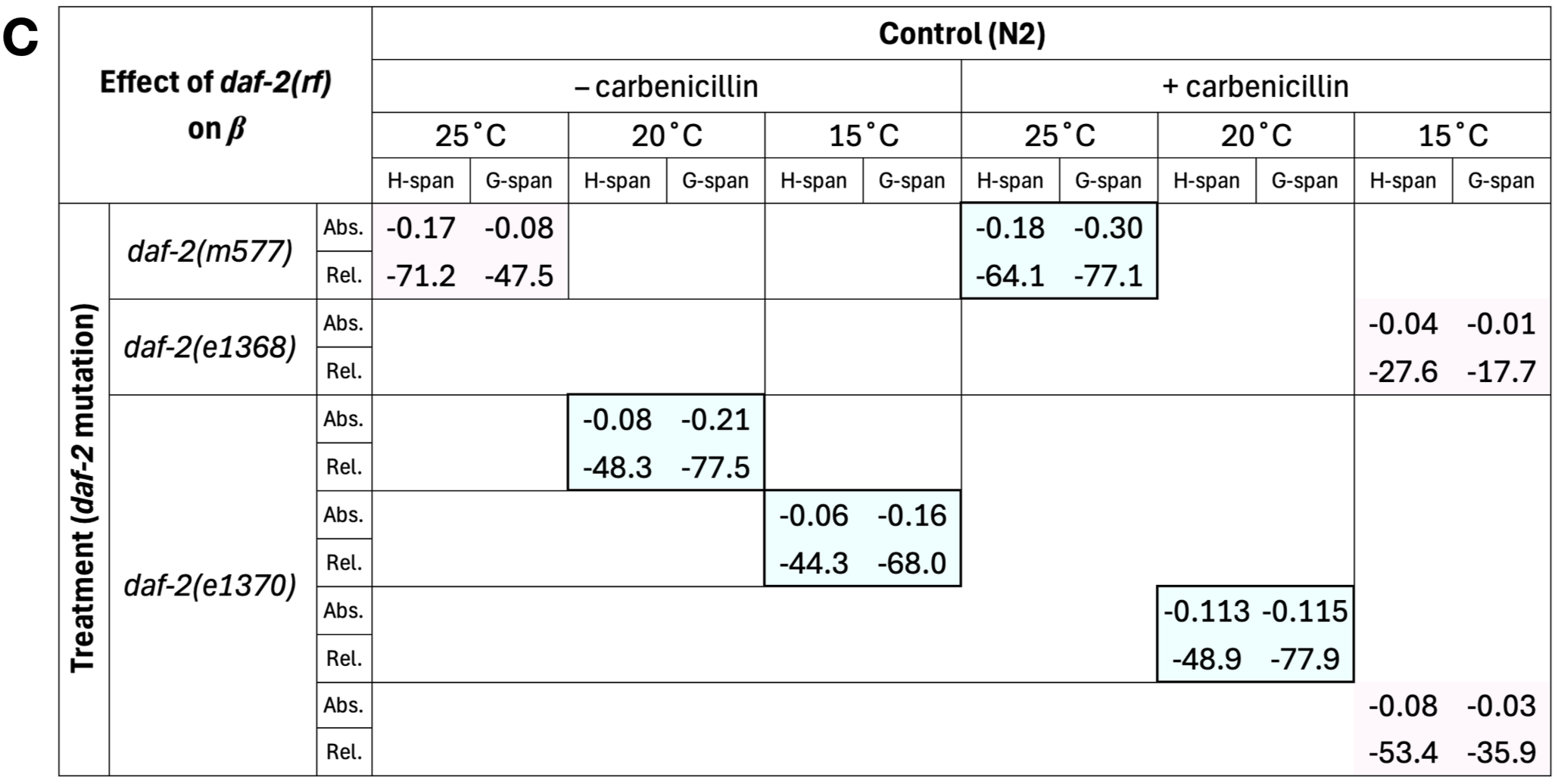

Table S2.

**Effect of changes in H-span and G-span on the *β* parameter.** All longevity treatments causing simultaneous *β* decrease and G-span^rel^ increase are presented, from three treatment categories: (**A**) low temperature, (**B**) carbenicillin, and (**C**) *daf-2(rf)*. N2, wild-type. Blue cells indicate treatments causing extended twilight longevity (ETL), and pink cells indicate treatments causing inverted ETL (see main text). Treatments where G-span causes greater reduction in *β* than H-span are enclosed by a bold border. Both absolute (Abs.) and relative percentage (Rel.) effects are shown, with the sign (positive/negative) indicating direction of change. For this analysis, *β* was obtained by maximum likelihood estimation in WinModest using H-span and G-span data (instead of lifespan data), for control and treatment cohorts, and the change (between control and treatment) quantified to estimate the separate contributions of H-span and G-span to lifespan *β* (H-span and G-span sum to lifespan and have additive effects on *β*).

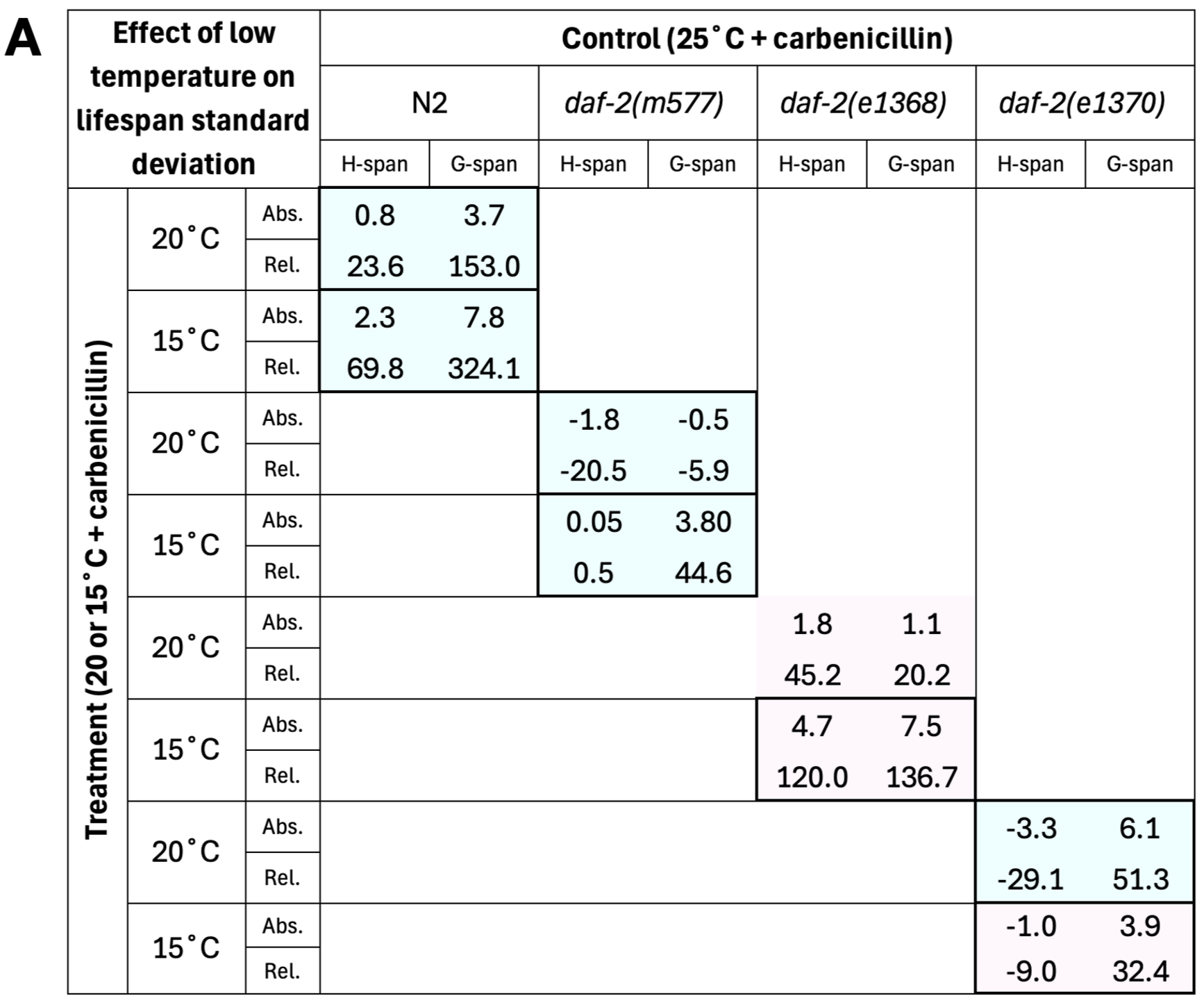

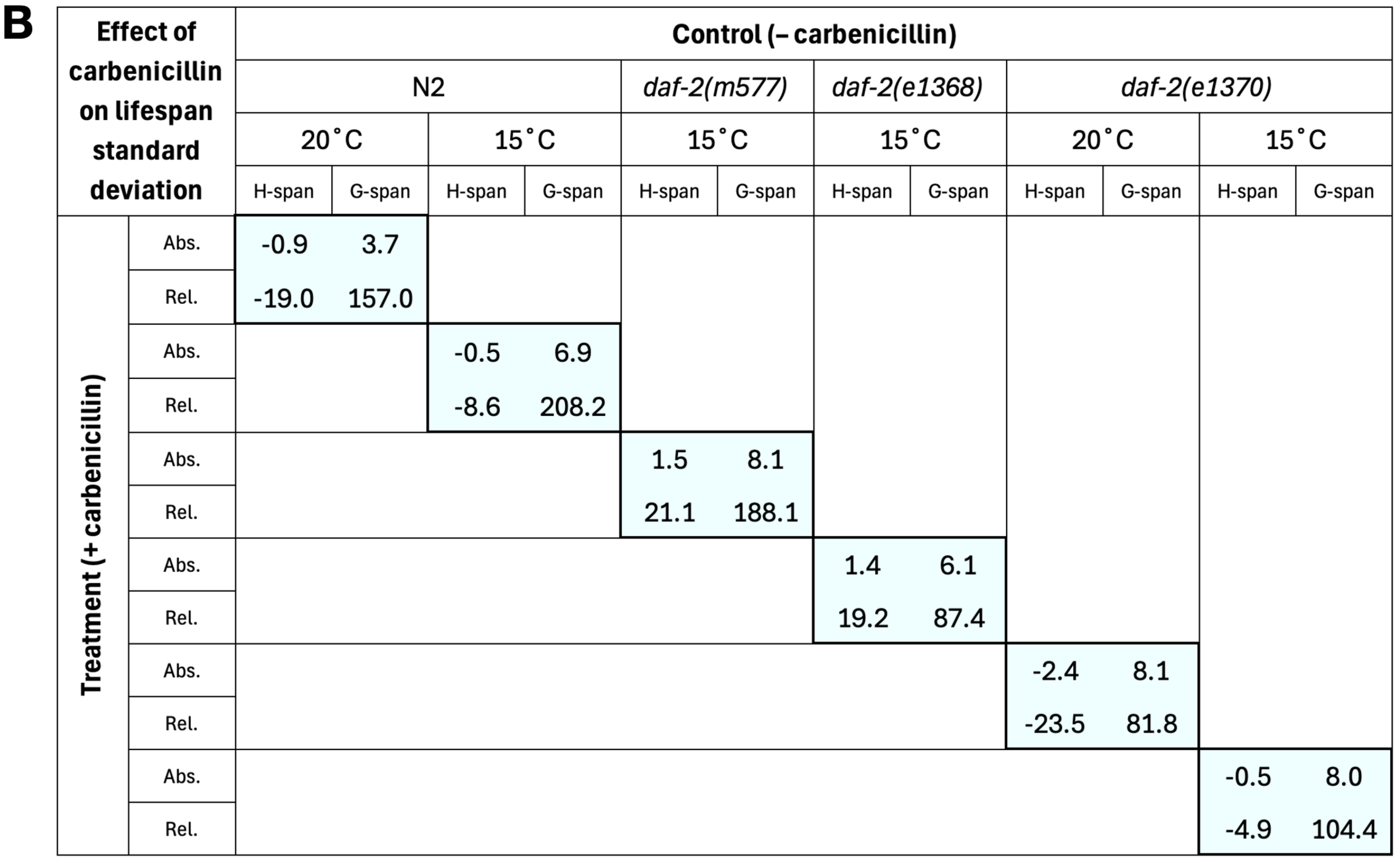

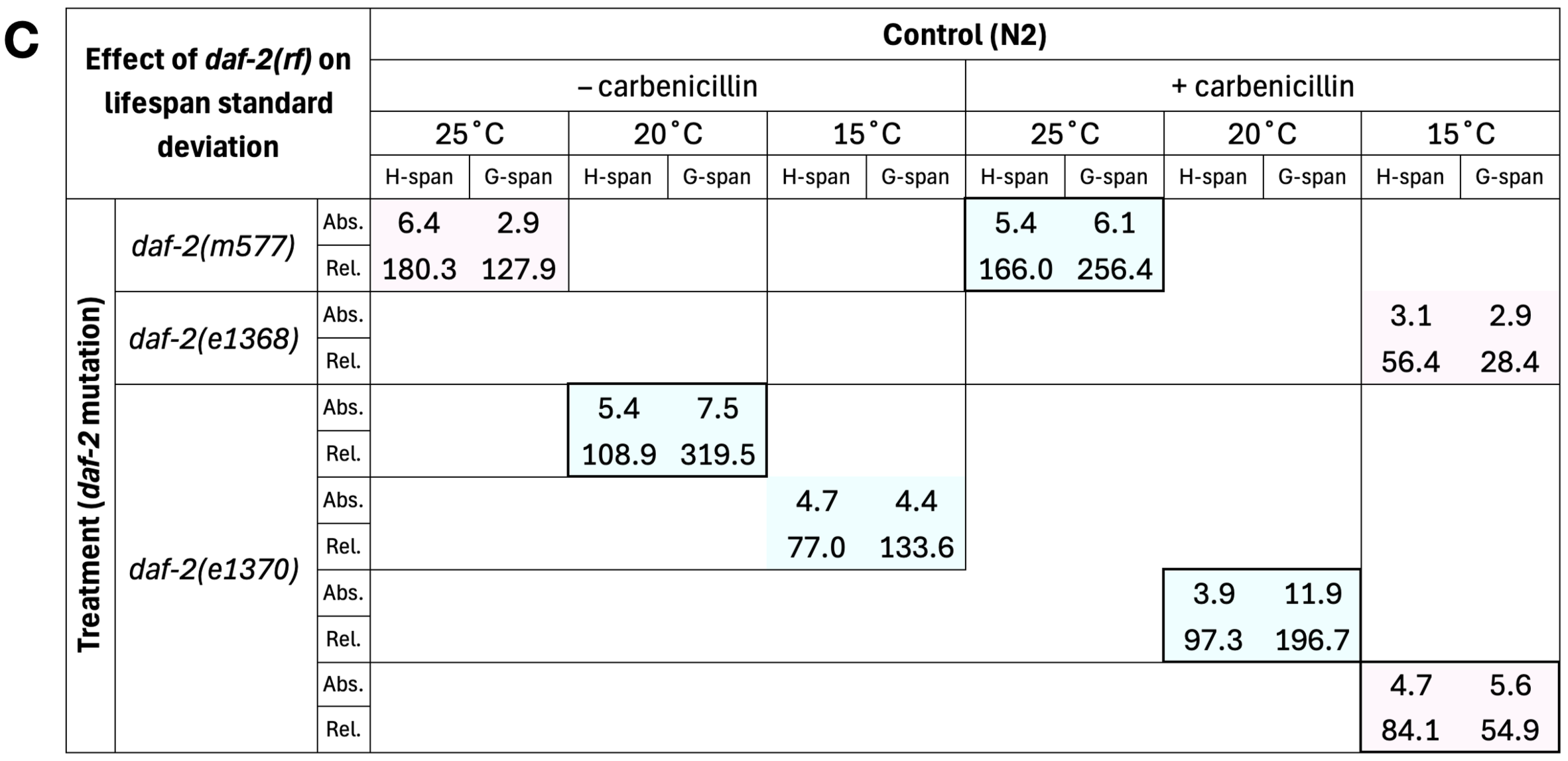

Table S3.
Effect of changes in H-span and G-span on lifespan standard deviation. All longevity treatments causing simultaneous *β* decrease and G-span^rel^ increase are presented, from three treatment categories: (A) low temperature, (B) carbenicillin, and (C) *daf-2(rf)*. N2, wild-type. Blue cells indicate treatments causing extended twilight longevity (ETL), and pink cells indicate treatments causing inverted ETL (see main text). Treatments where G-span causes greater increase in lifespan standard deviation than H-span are enclosed by a bold border. Both absolute (Abs.) and relative percentage (Rel.) effects are shown, with the sign (positive/negative) indicating direction of change (positive changes are related to *β* reduction). For this analysis, standard deviation was obtained for H-span and G-span, for control and treatment cohorts, and the change (between control and treatment) quantified to estimate the separate contributions of H-span and G-span to lifespan standard deviation (H-span and G-span sum to lifespan and have additive effects on lifespan standard deviation).

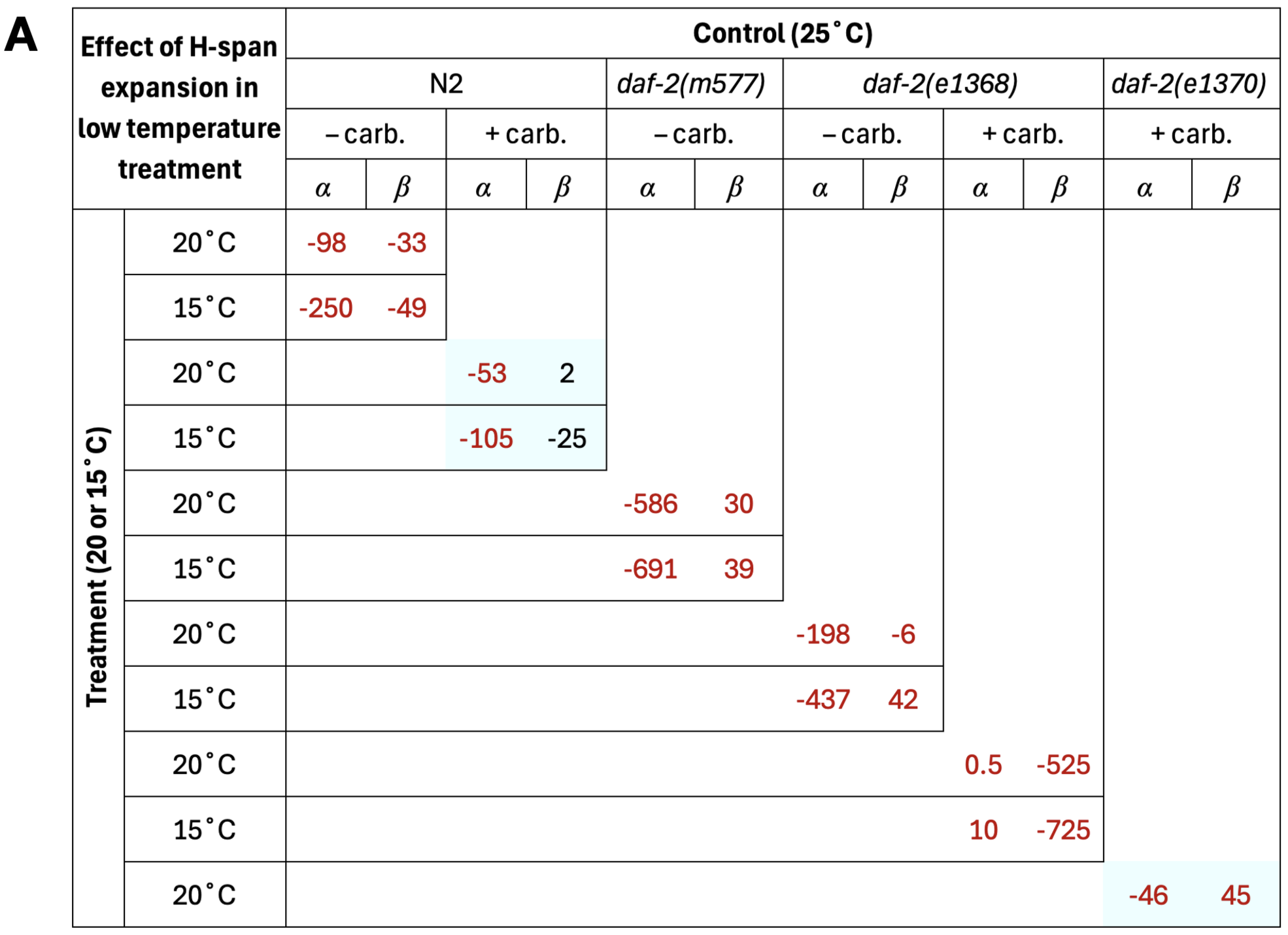

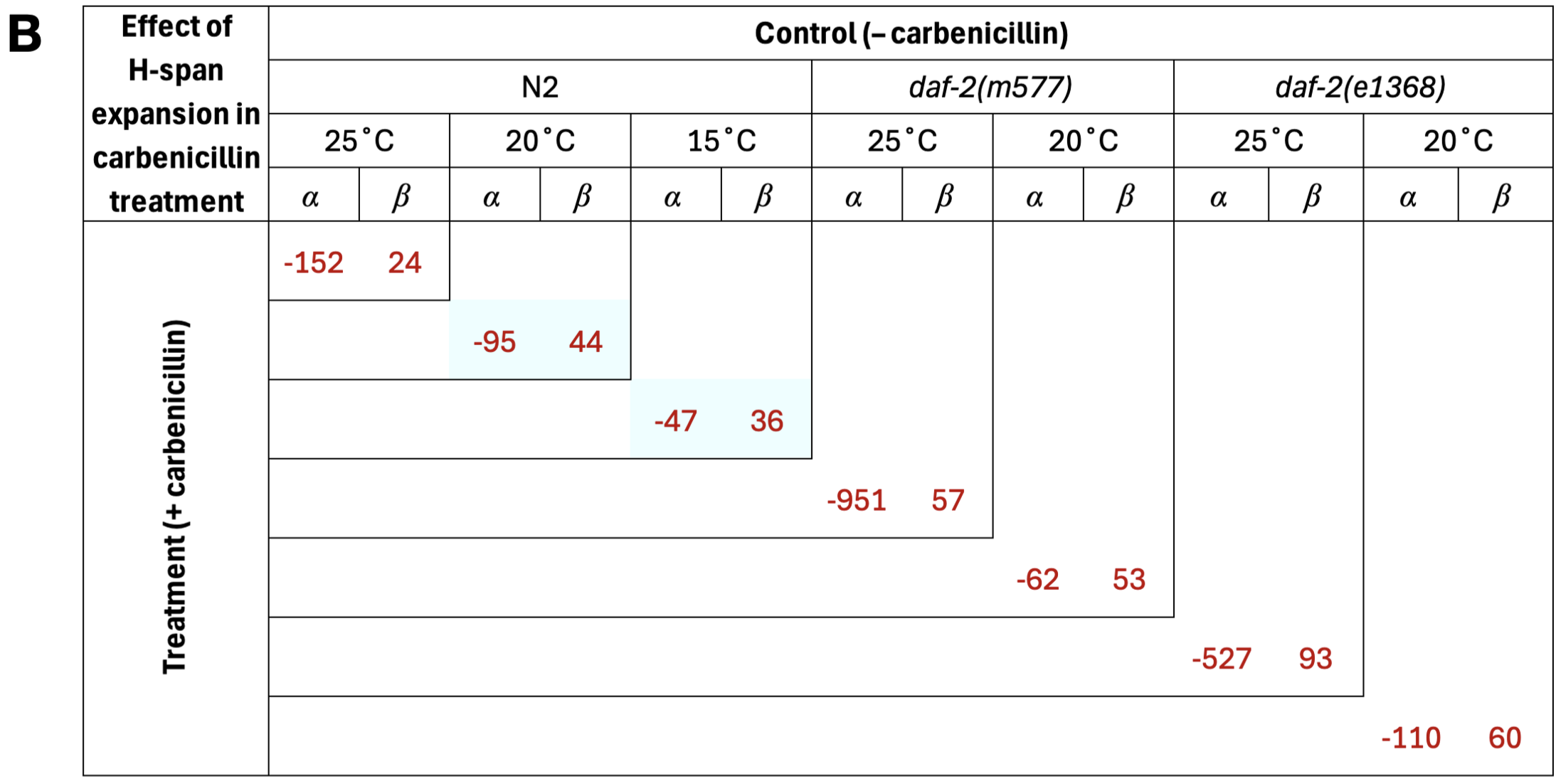

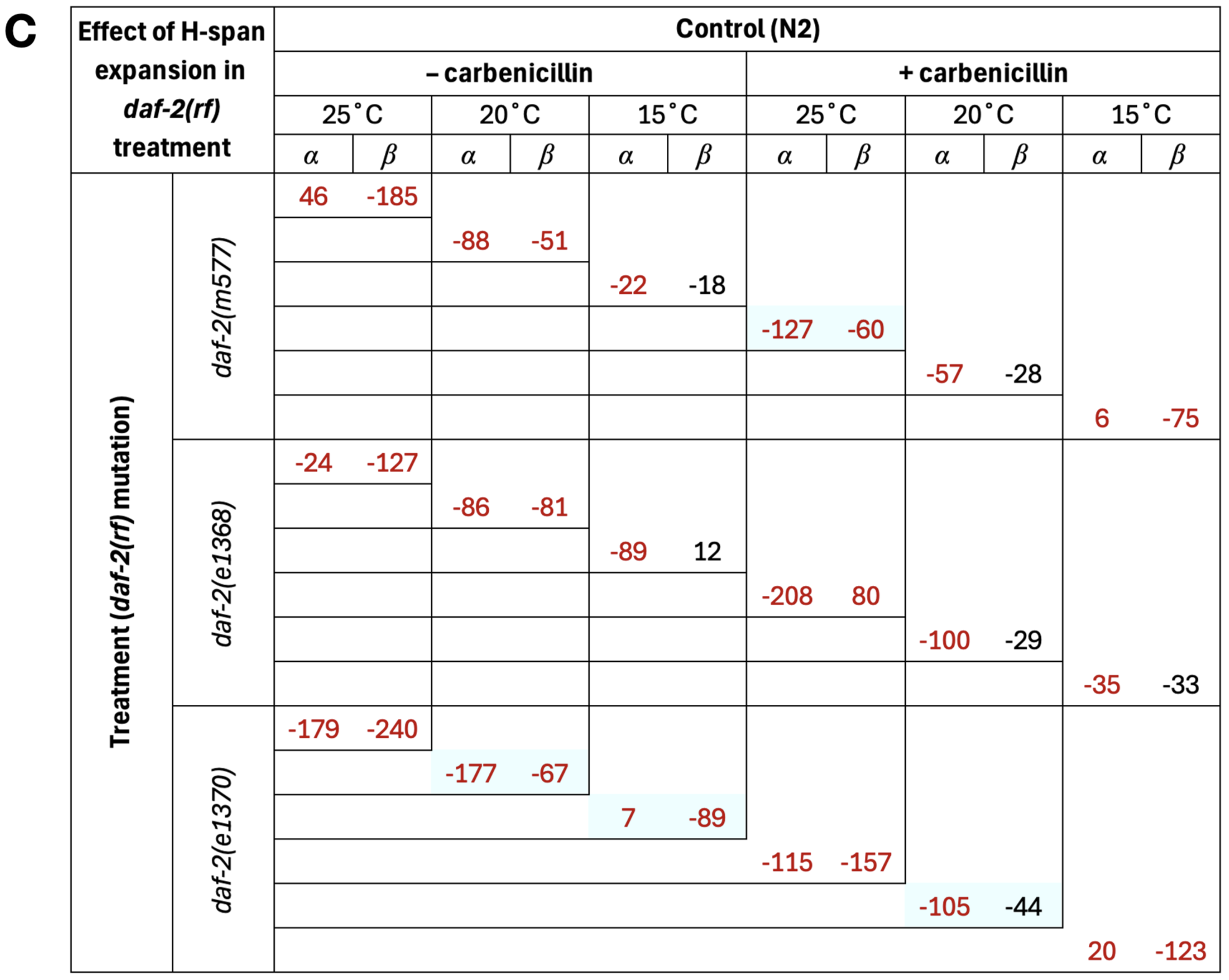

Table S4.
Effect of absolute H-span increase on *α* and *β*. All longevity treatments causing H-span^abs^ increase are presented, from three treatment categories: (A) low temperature, (B) carbenicillin, and (C) *daf-2(rf)*. N2, wild-type. Blue cells indicate treatments causing extended twilight longevity (ETL). Relative percentage effects are shown, with the sign (positive/negative) indicating direction of change, and statistically significant (*p*<0.05) effects are colored red. Effects of *α* and *β* were estimated as changes in the y-intercept and gradient of the least-squares linear regression model of H-span^abs^ over survival proportion (e.g. as plotted in Fig. 2B-G, upper panels). Increases in y-intercept and gradient correspond, respectively, to decreases in *α* and *β*. Statistical significance of these changes was assessed by two-tailed regression t-tests on model parameter estimates (y-intercept, treatment term; gradient, interaction term between treatment and survival proportion).

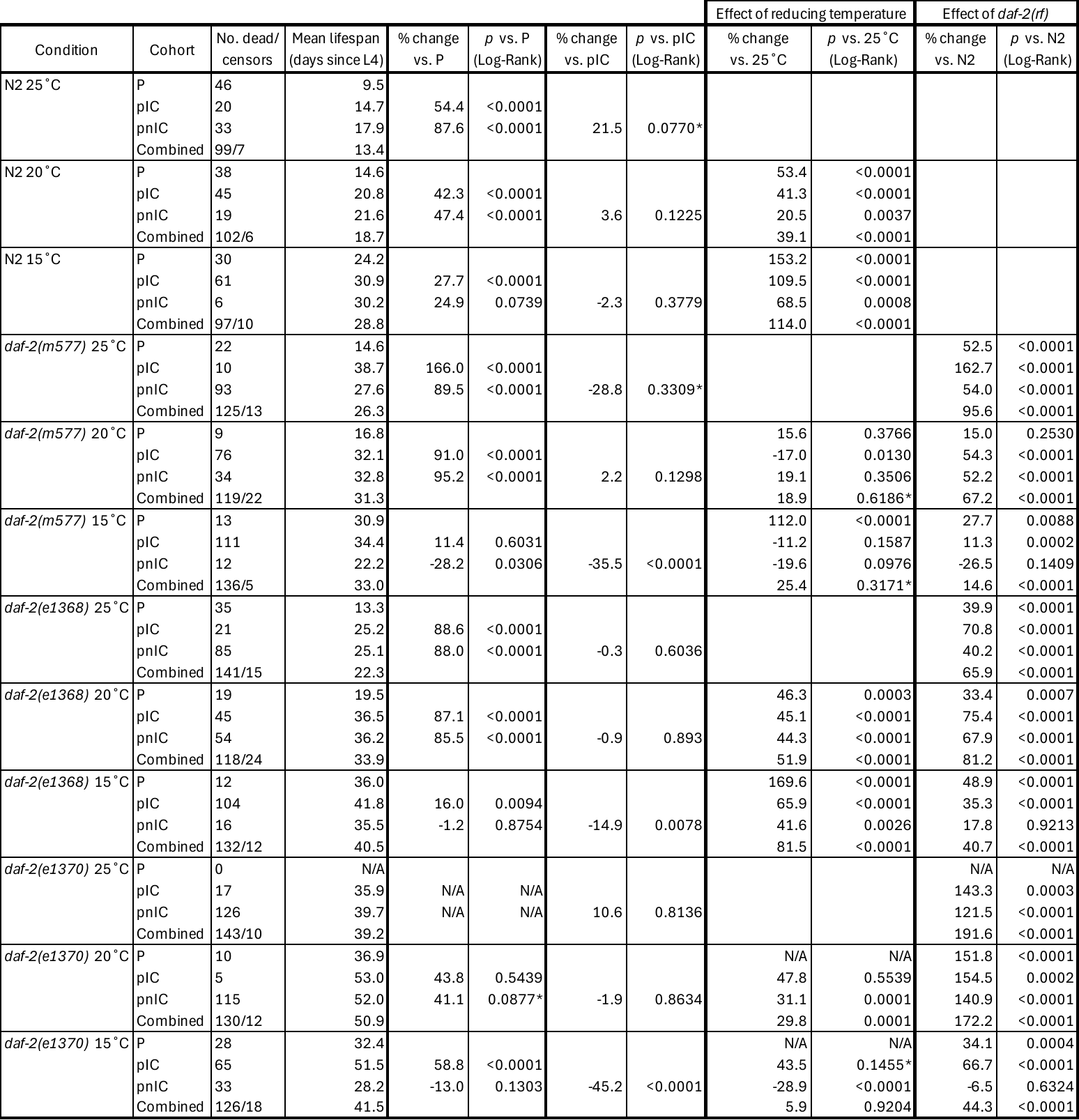

Table S5.
Lifespan data for P, pIC and pnIC subpopulations in non-carbenicillin cohorts, their differences and effect on them of low temperature and *daf-2(rf)* treatment. N2, wild-type. Mean lifespan was obtained by Kaplan-Meier analysis using age-specific mortality pseudofrequencies, to allow unbiased incorporation of censors in subpopulation analyses (see Methods). Mean lifespan of the subpopulations and combined population was obtained by Kaplan-Meier analysis using these mortality pseudofrequencies. *, cohorts where Log-Rank *p*>0.05, due to crossing survival curves and/or greater changes in early than late mortality; the Wilcoxon test is more appropriate to detect lifespan differences in these conditions (0.01<*p*<0.05 for inter-subpopulation comparisons, and 0.0002<*p*<0.006 for treatment effects; data not shown).

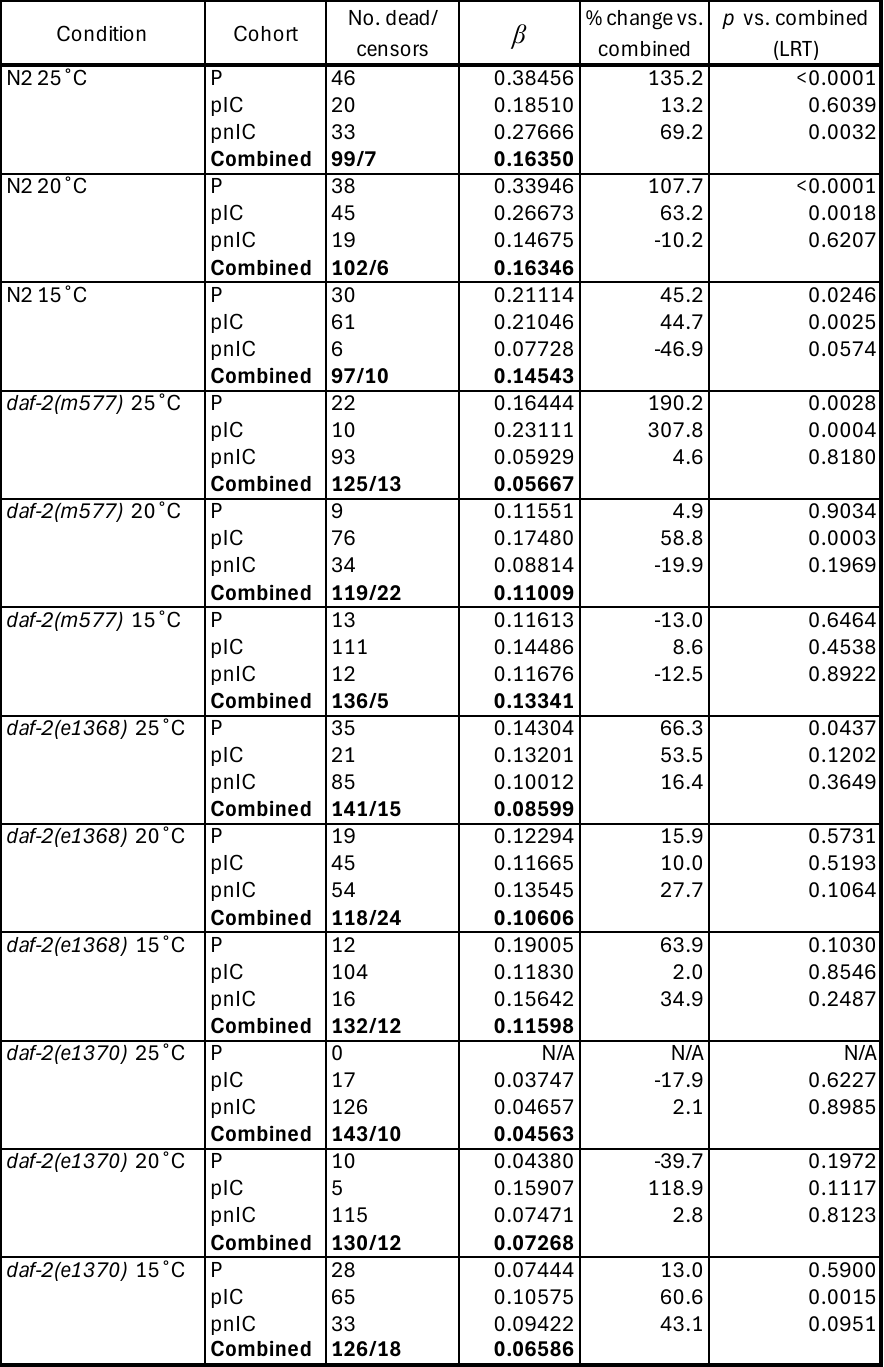

Table S6.
*β* parameters for P, pIC and pnIC subpopulations in non-carbenicillin cohorts, and their differences. N2, wild-type. *β* parameters were obtained by maximum likelihood estimation using mortality pseudofrequencies (in place of standard mortality frequencies) (see Methods). LRT, likelihood ratio test, used here to assess statistical significance of *β* differences between subpopulations and whole populations (combined).
